# Regulation of pSYSA defense plasmid copy number in *Synechocystis* through RNase E and a highly transcribed asRNA

**DOI:** 10.1101/2022.11.30.518505

**Authors:** Alena Kaltenbrunner, Viktoria Reimann, Ute A. Hoffmann, Tomohiro Aoyagi, Minori Sakata, Kaori Nimura-Matsune, Satoru Watanabe, Claudia Steglich, Annegret Wilde, Wolfgang R. Hess

## Abstract

Synthetic biology approaches toward the development of cyanobacterial producer strains require the availability of appropriate sets of plasmid vectors. A factor for the industrial usefulness of such strains is their robustness against pathogens, such as bacteriophages infecting cyanobacteria. Therefore, it is of great interest to understand the native plasmid replication systems and the CRISPR-Cas based defense mechanisms already present in cyanobacteria. In the model cyanobacterium *Synechocystis* sp. PCC 6803, four large and three smaller plasmids exist. The ∼100 kb plasmid pSYSA is specialized in defense functions by encoding all three CRISPR-Cas systems and several toxin-antitoxin systems. The expression of genes located on pSYSA depends on the plasmid copy number in the cell. The pSYSA copy number is positively correlated with the expression level of the endoribonuclease E. As molecular basis for this correlation we identified the RNase E-mediated cleavage within the pSYSA-encoded *ssr7036* transcript. Together with a cis-located abundant antisense RNA (asRNA1), this mechanism resembles the control of ColE1-type plasmid replication by two overlapping RNAs, RNA I and II. In the ColE1 mechanism, two non-coding RNAs interact, supported by the small protein Rop, which is encoded separately. In contrast, in pSYSA the similar-sized protein Ssr7036 is encoded within one of the interacting RNAs and it is this mRNA that likely primes pSYSA replication. Essential for plasmid replication is furthermore the downstream encoded protein Slr7037 featuring primase and helicase domains. Deletion of *slr7037* led to the integration of pSYSA into the chromosome or the other large plasmid pSYSX. Moreover, the presence of *slr7037* was required for successful replication of a pSYSA-derived vector in another model cyanobacterium, *Synechococcus elongatus* PCC 7942. Therefore, we annotated the protein encoded by *slr7037* as Cyanobacterial Rep protein A1 (CyRepA1). Our findings open new perspectives on the development of shuttle vectors for genetic engineering of cyanobacteria and of modulating the activity of the entire CRISPR-Cas apparatus in *Synechocystis* sp. PCC 6803.

## INTRODUCTION

The photosynthetic *Synechocystis* sp. PCC 6803 (from here: *Synechocystis* 6803) is a widely-used unicellular model cyanobacterium. The complete sequence of its chromosome was determined as early as 1996 (Kaneko et al., 1996) making it the first phototrophic and the third organism overall for which a major part of its genome sequence became available. In addition to its circular chromosome, the four large plasmids pSYSA, pSYSG, pSYSM, pSYSX (Kaneko et al., 2003) and three smaller plasmids pCC5.2, pCA2.4 and pCB2.4 (5.2 kb, 2.4 kb, and 2.3 kb) exist in *Synechocystis* 6803, which were all eventually sequenced as well (Yang and McFadden, 1993, 1994; Xu and McFadden, 1997). Although the combined coding capacity of these seven plasmids makes up almost 10% of all protein-coding genes in *Synechocystis* 6803, their biological functions have only partially been addressed. Recently, a large gene cluster for the biosynthesis of an extracellular sulfated polysaccharide, called synechan, was identified on the ∼120 kb plasmid pSYSM (Maeda et al., 2021).

Of particular interest is the plasmid pSYSA. This plasmid was previously characterized as specialized for defense functions by encoding all three CRISPR-Cas systems in *Synechocystis* 6803 (Scholz et al., 2013; Reimann et al., 2017; Behler et al., 2018) and at least seven distinct type II toxin-antitoxin systems (Kopfmann and Hess, 2013; Kopfmann et al., 2016). Resequencing of the *Synechocystis* 6803 lab strain PCC-M in 2012 revealed two deletion events in pSYSA (Trautmann et al., 2012) compared to the original sequence determined 10 years before (Kaneko et al., 2003). Both of these deletions were mapped to the CRISPR-Cas system, removing 2,399 bp and 159 bp, respectively (Trautmann et al., 2012).

While the small endogenous plasmids in *Synechocystis* 6803 replicate by the rolling circle mechanism (Yang and McFadden, 1993, 1994; Xu and McFadden, 1997), the replication type of the large plasmids and also of plasmids in other cyanobacteria have largely remained unknown so far.

Previous analyses after the transient inactivation of RNase E by temperature shift (TIER-seq) revealed that transcripts derived from the four major plasmids accumulated differentially between cells expressing an unmodified form of this riboendonuclease, *rne*(WT) and a temperature-sensitive form, *rne*(Ts) (Hoffmann et al., 2021). Moreover, the joint overexpression of RNase E and RNase HII led up to 3.8- and 2.4-fold increased copy numbers of plasmids pSYSA and pSYSM, respectively (Hoffmann et al., in progress). These results were interpreted as pointing toward a possible role of RNase E in copy number regulation of these replicons. Indeed, such functions were described for certain endoribonucleases. RNase E is involved in the control of ColE1-type plasmid replication in *E. coli* (Lin-Chao and Cohen, 1991) and RNases J1 and J2 in the RepA_N-family plasmid pSA564 replication in *Staphylococcus aureus* (Guimarães et al., 2021).

Here we show that two abundant and overlapping pSYSA transcripts, the *ssr7036* mRNA and the asRNA1, are substrates for RNase E *in vivo* and *in vitro*. Manipulation of their levels and processing patterns leads to an altered plasmid copy number. Transfer of this locus to an *E. coli* pUC vector yielded a plasmid capable of self-replication in *Synechocystis* 6803 and, if the Slr7037 replication protein was present, also in the unrelated *Synechococcus elongatus* PCC 7942 (from here *Synechococcus* 7942). In addition, deletion of *slr7037* led to the integration of pSYSA into the chromosome or the other large plasmid pSYSX, suggesting that Slr7037 is crucial for pSYSA replication. This system resembles the copy control mechanism of ColE1-type plasmids, but differs upon closer inspection in several details.

## MATERIALS AND METHODS

### Culture media, strains, growth conditions and manipulation of cyanobacteria

If not stated otherwise, the *Synechocystis* 6803 strains PCC-M was used as wild type as re-sequenced in 2012 (Trautmann et al., 2012). Cultures were grown in BG-11 (Rippka et al., 1979) substituted with 0.3% (w/v) sodium thiosulfate and 10 mM N-[Tris- (hydroxy-methyl)-methyl]-2-aminoethanesulfonic acid (TES) buffer (pH 8.0). Liquid cultures were grown in Erlenmeyer flasks at 30°C under continuous white light illumination (30 µmol photons m^-2^ s^-1^ or as stated) and with constant shaking at 135 rpm. Plate cultures were grown on 0.75% bacto-agar BG-11 plates with added antibiotics as needed (kanamycin, 50 µg/mL, chloramphenicol, 10 µg/mL, gentamicin, 2 µg/mL).

*Synechocystis* 6803 was transformed with the plasmids VIII.22, VIII.23, VIII.44 and VIII.45 (**Table 1**). For one transformation, aliquots corresponding to 20 OD_750_ units were taken from exponentially growing cultures at an OD_750_ between 0.6 and 0.9. Cells were collected by centrifugation at 3,237 g for 6 min in a swing-out rotor at room temperature and resuspended in a droplet of remaining BG-11. Five µg of plasmid DNA were added. Thereafter, cells were incubated for 1.5 h in the light and then plated on 0.9% bacto-agar BG-11 plates. After 24 h, 400 µL of BG-11 supplemented with 50 µg/mL kanamycin (calculated according to the total plate volume) were pipetted below the agar. After 14 days at 30°C with a light intensity of 50 μmol photons m^−2^ s ^−1^, first colonies appeared. These were transferred onto fresh 0.75% bacto-agar BG-11 plates containing 40 µg/mL kanamycin.

**Table 1.** Strains, plasmid vectors and used constructs.

| Strain | Description | Reference |
| --- | --- | --- |
| *E. coli* DH5α | F<sup>-</sup> ϕ80/*lacZ*ΔM15 Δ(*lacZYA-argF*)U169 *recA1 endA1 hsdR17*(r<sub>K</sub><sup>-</sup>, m<sub>K</sub><sup>+</sup>) *phoA supE44 λ-thi-1 gyrA96 relA1* | Thermo Fisher Scientific |
| *E. coli* Top10F' | identical to TOP10 cells, with the addition of an F' episome | Thermo Fisher Scientific |
| *E. coli* M15 [pREP4] pQE70-*rne* | Expression of recombinant *Synechocystis* 6803 RNase E | (Behler et al., 2018) |
| *Synechocystis* 6803 pVZΔKm<sup>R</sup> | *slr1128::kanR 3xFLAG-rne<sup>+</sup>* pVZ321Δ(9220-1203) | (Hoffmann et al., 2021) |
| *Synechocystis* 6803 3xFLAG-*rne* | *slr1128::kanR 3xFLAG-rne<sup>+</sup>* | (Hoffmann et al., 2021) |
| *Synechocystis* 6803 *rne*(WT) | Δ(*rne-rnhB*) *rnep::kanR* pVZ321Δ(9,220-1,203) (9,219)::(*rnep-3xFLAG-rne<sup>+</sup>-rnhB-rnhBt*) | (Hoffmann et al., 2021) |
| *Synechocystis* 6803 *rne*(5p) | Δ(*rne-rnhB*) *rnep::kanR* pVZ321Δ(9,220-1,203) (9,219)::(*rnep-3xFLAG-rne*(5p)-*rnhB-rnhBt*) | (Hoffmann et al., in progress) |
| *Synechocystis* 6803 Δ*slr7037-1* | Gene *slr7037* deleted by replacement with a chloramphenicol resistance cassette (pSYSA-chromosome integration via *slr7104-slr7105/sll0699-sll0700*) | This work |
| *Synechocystis* 6803 Δ*slr7037-2* | Gene *slr7037* deleted by replacement with a chloramphenicol resistance cassette (pSYSA-chromosome integration via *slr7104-slr7105/sll1256-sll1257*) | This work |
| *Synechocystis* 6803 Δ*slr7037-3* | Gene *slr7037* deleted by replacement with a chloramphenicol resistance cassette (pSYSA-pSYSX integration via *slr7005/sll6109*) | This work |
| Plasmid | Description | Reference |
| pVZ322 | IncQ KmR GmR Mob<sup>+</sup>; replicative vector for cyanobacteria | (Zinchenko et al., 1999) |
| VI.16 | renamed here from pVZ322 derivative (pVZ322s) in which kanamycin resistance marker was deleted | (Behler et al., 2018) |
| VIII.17 | pVZ322s::P<sub>rha</sub>-7036-3xFLAG | This work |
| VIII.18 | pVZ322s::P<sub>rha</sub>-GFP-3xFLAG | This work |
| VIII.22 | pUC19s with inserted *ssr7036* and kanamycin resistance genes | This work |
| VIII.23 | pUC19s backbone and the entire *ssr7036-slr7037* region including the pSYSA origin of replication | This work |
| VIII.44 | pUC19s::nP7036-7037_P<sub>petE</sub>sfGFP_KmR (VIII.23 plus sfGFP under control of P<sub>petE</sub>) | This work |
| VIII.45 | pUC19s::nP7036-7037STOP_KmR | This work |
| VII.48 | pUC19::T1-P<sub>rha</sub>-RBS-eYFP-3xFLAG-T2-KmR-RhaS-T3 amplified from pCK355 | (Kelly et al., 2019); Luisa Hemm, unpublished |
| VIII.58 | pVZ322s::P<sub>rha</sub>-7036STOP-3xFLAG (VIII.17 with 2<sup>nd</sup> codon modified to STOP codon in *ssr7036*) | This work |
| pUC19 | GenBank entry L09137.2 | (Yanisch-Perron et al., 1985) |
| pUC19s | pUC19 backbone of 1,965 bp lacking nt 26 to 746 from GenBank entry L09137.2 | This work |

The pVZ322 (Zinchenko et al., 1999) based plasmids VI.16, VIII.17, VIII.18 and VIII.58 were transferred into the *Synechocystis* cells by electroporation. Twenty-five mL of wild-type culture with an OD_750_ of 0.8 – 1.0 were harvested at 3,237 g for 10 min at room temperature in a swing-out rotor. The cell pellet was resuspended in 2 mL ice-cold HEPES (1 mM, pH 7.5) and the cells were collected as before. This wash step was repeated twice. Thereafter, the pellet was resuspended in 100 µL ice-cold HEPES per approach and 1 µg of plasmid DNA was added. The electrocompetent cells were transferred into ice-cold electroporation cuvettes. The electroporation was performed at 2,500 V for 4 ms in an electropulser (MicroPulser, Bio-Rad). Electroporated cells were resuspended first in 1 mL BG-11 medium, then added to 50 mL BG-11 in an Erlenmeyer flask and incubated for 24 h at 30°C and 50 µmol photons m^-2^ s^-1^. Thereafter, cells were harvested at 3,237 g for 10 min, resuspended in a drop of the supernatant and spread onto 0.75% bacto-agar BG-11 plates supplemented with 2 µg/mL gentamycin. Colonies began to appear after eight days.

For transformation of *Synechococcus* 7942, a 2 mL culture at an OD_750_ of 1.0 to 1.7 was harvested by centrifugation at 11,000 *g* for 2 min at room temperature. The supernatant was removed and the pellet resuspended in 1 mL fresh BG-11. The cells were centrifuged again at 11,000 *g* for 1 min at room temperature, the supernatant removed and the cells resuspended in 100 µL BG-11 by gently pipetting up and down before 5 µg of plasmid DNA were added. This mixture was incubated at 34°C for 4 h in the dark, then plated on 0.9% bacto-agar BG-11 plates. The following addition of kanamycin and the selection conditions were identical to the procedure described for the transformation of *Synechocystis* 6803. After 8 days first transformed colonies appeared.

### Construction of mutant strains

All vectors and constructs used in this study are listed in **Table 1**. Construction of strains *rne*(WT), *rne*(Ts) and *rne*(5p) were previously described (Hoffmann et al., 2021). For the construction of plasmids VIII.22 and VIII.23, the primers P1 and P2 were used to inversely amplify the pUC19 backbone (for primer details see **Table 2**). The gene *ssr7036* including its 5’ and 3’ untranslated regions (UTRs) was amplified using the primers P3 and P4 and *Synechocystis* 6803 genomic DNA as template. Genes *ssr7036* and *slr7037* including the intergenic region and their 5’ and 3’ UTRs were amplified with primers P3 and P5. The plasmids were constructed using AQUA cloning (Beyer et al., 2015) and chemocompetent *E. coli* DH5α cells (**Table 1**). The plasmid VIII.44 was constructed using primers P2 and P6 to amplify the VIII.23 plasmid as backbone, primers P7 and P8 to amplify the *petE* promoter (P*_petE_*) and primers P9 and P10 to amplify the *sfGFP* gene from plasmid VII.65. The plasmid VIII.45 is identical to VIII.23 except for a point mutation in *slr7037* leading to an early stop codon. This point mutation was inserted through site-directed mutagenesis, using the primers P11 and P12 with the VIII.23 plasmid as a template.

**Table 2.**
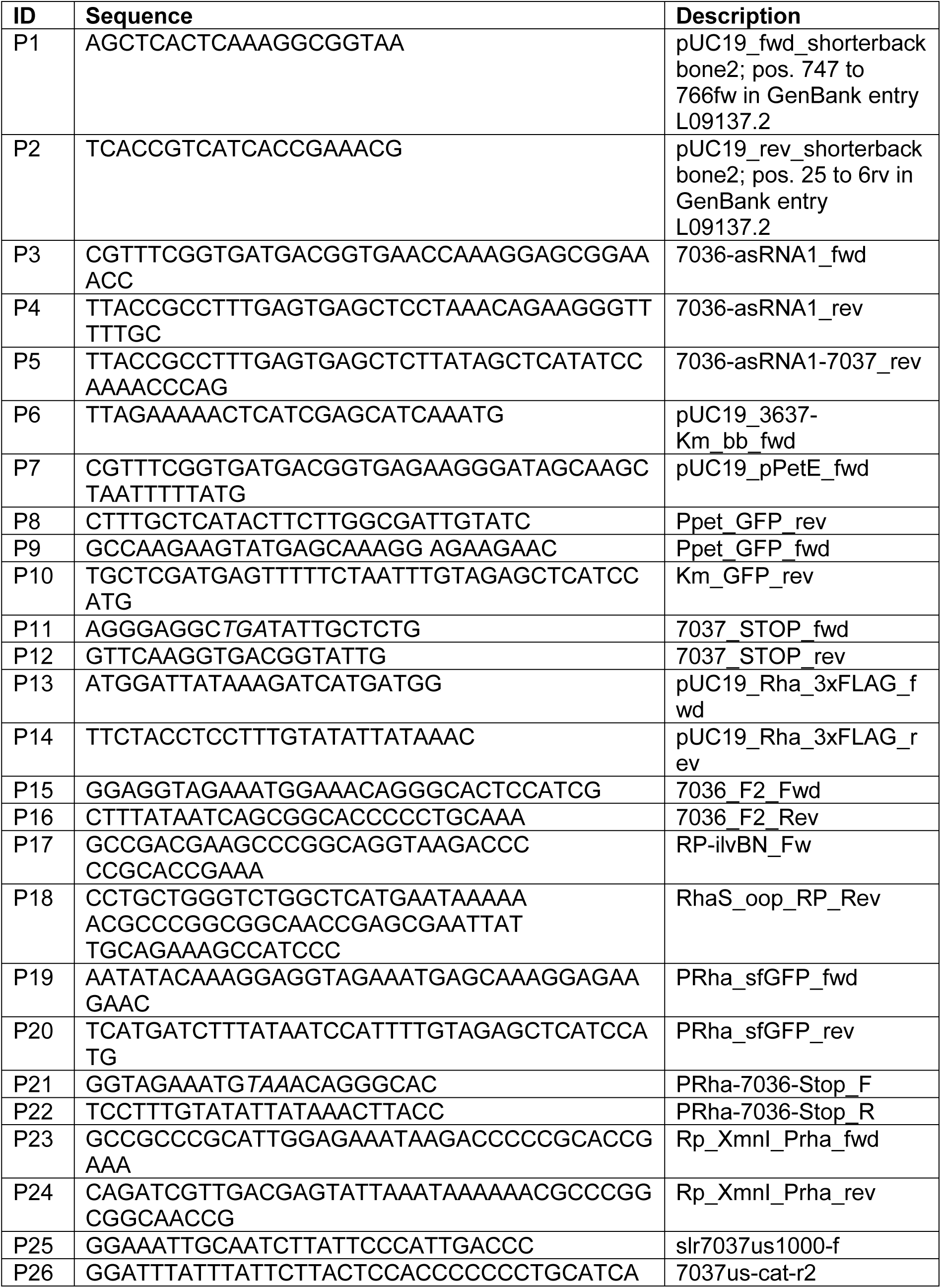
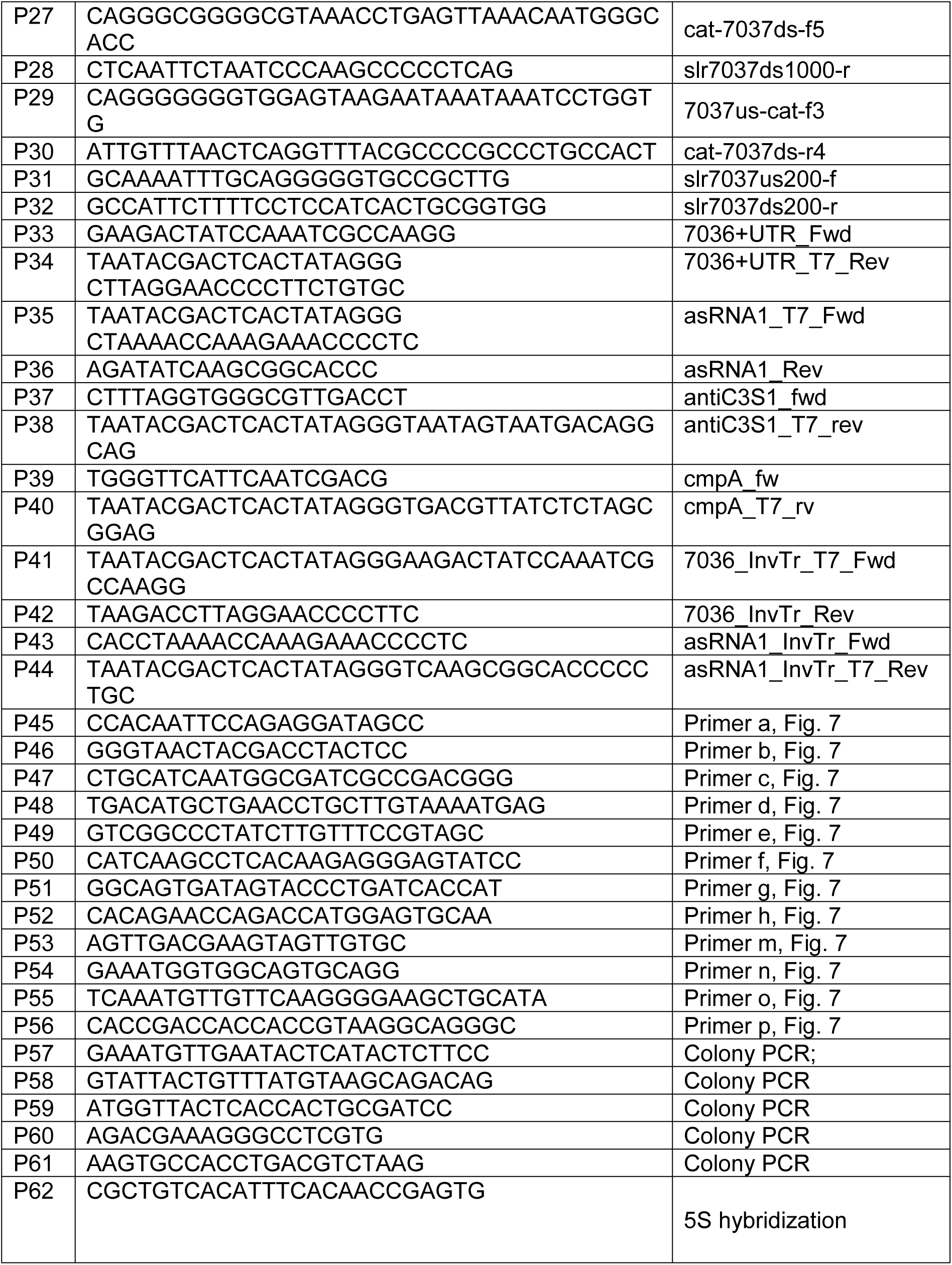
Oligonucleotides used in this study. The stop codon introduced with P11 is in italics.

Plasmid VIII.17 is based on plasmid VI.16 with the following insert: T1-P_Rha_-RBS-eYFP-3xFLAG-T2-*KmR*-RhaS-T3 that was amplified from VII.48. T1-T3 designate the *ilvBN*, ECK120034435 (http://parts.igem.org/Part:BBa_K2243023) and bacteriophage λ OOP rho-independent transcriptional terminators (Dühring et al., 2006; Kelly et al., 2019). The plasmid VII.48 was inverse PCR amplified excluding eYFP, using primers P13 and P14. The gene *ssr7036* was PCR amplified from genomic DNA using the primers P15 and P16. Both fragments were assembled and subcloned in *E. coli* Top10F’ yielding plasmid VIII.24 (**Table 1**). Using primers P17 and P18, the region T1-P_Rha_-RBS*-ss7036*-3xFLAG-T2-KmR-RhaS-T3 was PCR amplified with overhangs into a shortened version of pVZ322, called pVZ322s (**Table 1**). The backbone was digested with XmnI and both fragments were assembled and subcloned in *E. coli* Top10F’ yielding plasmid VIII.17. The plasmid VIII.18 was constructed the same way as VIII.17 but using primers P19 and P20 to amplify the sfGFP gene from plasmid VII.65.

Starting with plasmid VIII.17, the second codon of the *ssr7036* gene was modified to a stop codon (resulting in plasmid VIII.58) through site-directed mutagenesis using primers P21 and P22 and inverse PCR with plasmid VIII.24 as a template. Competent Top10F’ *E. coli* cells were transformed with the resulting PCR product. The plasmid DNA was then isolated and used as a template for PCR amplification of the mutagenized insert using primers P23 and P24. Chemocompetent Top10F’ *E. coli* cells were transformed with the PCR product and XmnI digested pVZ322s plasmid as backbone.

The PCR products were assembled with AQUA cloning (Beyer et al., 2015) in chemocompetent *E. coli* DH5α cells, if not stated otherwise. The PCRBio HiFi DNA polymerase was used in all PCRs throughout.

To construct DNA fragments for the deletion of *slr7037*, the regions flanking *slr7037* were amplified by PCR using primers P25 and P26 (for the upstream fragment) and P27 and P28 (for the downstream fragment), respectively. The chloramphenicol resistance marker gene was amplified from pUC303 (Kuhlemeier et al., 1983) using primers P29 and P30. Three fragments were combined by PCR using primers P25 and P28. All constructs were verified by sequencing. After transformation of DNA fragments into *Synechocystis* GT-I strain (Kanesaki et al., 2012), several clones showing resistance to chloramphenicol were isolated. The complete disruption of *slr7037* in strain Δ*slr7037* was confirmed by PCR using primers P31 and P32 with genomic DNA as a template. Three clones with complete deletion of *slr7037* were selected and used for further analyses (**Figure S1**).

### Preparation of total RNA and Northern blot hybridizations

25 to 50 mL of densely grown cultures were harvested at 3,237 *g* for 10 min at room temperature in a swing-out centrifuge and resuspended in 1 mL PGTX (Pinto et al., 2009). For cell lysis, the samples were incubated for 15 min at 65°C with occasional vortexing, followed by addition of 700 µL chloroform/isoamyl alcohol (24:1), mixing and incubation for 10 min at room temperature. Following a second chloroform extraction, the upper phase was precipitated with 1 vol of isopropanol overnight at -20°C. Pellets were collected by centrifugation at 13,226 *g* for 30 min at 4°C and washed once with 1 mL of 70% EtOH. The air-dried pellet was dissolved in 20 to 50 µL of RNase-free H_2_O.

To inactivate the temperature sensitive RNase E (*rne*(TS), (Hoffmann et al., 2021)) strains were incubated for 0 h, 1 h or 24 h at 39°C. Wildtype (WT) and a strain carrying a plasmid-encoded RNase E control (*rne*(WT), resulting in higher RNase E levels than in WT) were treated the same way. For Northern blot analysis, 10 µg samples of total RNA were separated by 8 M urea 10% PAGE and electroblotted on Hybond N+ nylon membranes (Cytiva) with 1 mA per cm^2^ for 1 h. Northern hybridizations were performed with radioactively labelled probes (primers P33 to P36 for templates) generated using [α-^32^P]-UTP and the Maxiscript T7 *in vitro* transcription kit (Thermo Fisher Scientific). After blotting the RNA was crosslinked to the membrane with 240 mJ using a UV-Stratalinker (Stratagene). Hybridizations were performed in Northern buffer (50% deionized formamide, 7% SDS, 250 mM NaCl and 120 mM Na_2_HPO_4_/NaH_2_PO_4_ pH 7.2) overnight at 62°C. The membranes were washed at 57°C in buffer 1 (2×SSC (3 M NaCl, 0.3 M sodium citrate, pH 7.0), 1% SDS), buffer 2 (1×SSC, 0.5% SDS) and buffer 3 (0.1×SSC, 0.1% SDS) for 10 min each. Signals were detected with the biomolecular imager Typhoon FLA 9500 (GE Healthcare) using phosphorimaging and a photomultiplier value of 1000 V.

### Western blot analysis

Cultures were induced with 1.25 µM Cu_2_SO_4_ at an OD_750_ between 0.6 and 1.0. After 24 h, the cultures were collected at 3,237 *g* for 10 min, the pellet was resuspended in 200 µL TBS and transferred into a 1.5 mL screw cap tube containing 150 µL glass beads (Retsch) with diameters of 0.1 and 0.25 mm. Cell disruption was carried out with the Precellys 24 homogenizer (Bertin Technologies) at 6,000 rpm for 3 repeats of 3 x 10 s with 5 s of break in between at 4°C. After that, the mixture was transferred into a Micro Bio Spin column in an 1.5 mL Eppendorf tube and centrifuged for 2 min at 100 g at 4°C to separate the glass beads from the sample. The flow through was centrifuged at 15,871 *g* for 30 min at 4°C to remove the cell debris. The supernatant was transferred into a fresh tube and protein concentration was measured using the Invitrogen Qubit 3 fluorometer (Thermo Fisher Scientific). Ten µg total protein were loaded per lane.

5x protein loading buffer (250 mM Tris, pH 6.8, 25% glycerol, 10% SDS, 500 mM DTT, 0.05% bromophenol blue) was added to the samples before denaturation at 95°C for 5 min. As marker the PageRuler™ Prestained Protein Ladder (Thermo Fisher Scientific) was used. Samples were run on a 6% SDS polyacrylamide stacking gel and a 10% SDS polyacrylamide separating gel to separate the proteins by size. The gel was blotted onto a nitrocellulose membrane (Hybond™-ECL, Cytiva) at 1.2 mA/cm^2^ for 1 h. The membrane was stained using Ponceau solution (0.2% Ponceau, 3% TCA) for 5 min shaking at 180 rpm at room temperature. To remove the Ponceau stain, the membrane was washed with deionized water. After the Ponceau staining, the membrane was washed 3 times for 5 min in TBS-T and blocked overnight in TBS-T with 3% milk powder at 4°C. The next day, the membrane was washed again 3 times for 10 min in TBS-T, then incubated for 1 h at room temperature in 20 mL TBS-T with 3% milk powder and 1:5,000 diluted anti-GFP antiserum (Abcam) while shaking with 180 rpm. Subsequently, the membrane was washed three times for 10 minutes in TBS-T, then incubated for 1 h in 20 mL TBS-T with 3% milk powder and 1:10,000 diluted goat anti-rabbit IgG-HRP (Sigma). Finally, the membrane was washed for 10 min twice in TBS-T and briefly once in TBS. The membrane was sprayed with WesternBright ECL Spray (Advansta) and the signal was visualized using the FUSION SL Transilluminator (Vilber Lourmat).

### Total DNA preparation and Southern blot hybridization

Total DNA was prepared from *Synechocystis* 6803 by collecting cells from 50 mL cultures by centrifugation at 3,237 *g* for 10 min. The pellets were resuspended in 1 mL SET buffer (50 mM Tris, pH 7.5; 1 mM EDTA, 25% (w/v) sucrose). For lysis, resuspended cells were incubated in 100 mM EDTA (pH 8), 2% SDS (w/v) and 100 µg/mL proteinase K at 50°C for 16 h. DNA was extracted by phenol/chloroform/isoamylalcohol (25:24:1) extraction. DNA was precipitated from the final aqueous phase by adding 1 vol isopropanol, incubation at -20°C for at least 2 h and centrifugation at 15,871 *g* for 30 min at 4°C. The pellet was washed with 70% ethanol, air-dried and resuspended in 30 µL sterile Milli-Q water.

For restriction analysis, 4 µg of total DNA each were digested with FastDigest HindIII restriction endonuclease for 3 h at 37°C to ensure complete digestion. The digested DNA samples were subjected to gelelectrophoretic separation for 1 to 2 h at room temperature on ethidium bromide stained 0.8% agarose gels with 120 V. Thereafter, the gels were incubated at 70 rpm at room temperature for 30 min each in 0.25 M HCl, in denaturation solution (1.5 M NaCl, 0.5 M NaOH) and in neutralization solution (1.5 M NaCl, 0.5 M Tris pH 7.5) prior to blotting. The DNA samples were blotted onto Hybond-N+ nylon membranes (Cytiva) by capillary transfer overnight with 20x SSC (3 M NaCl, 300 mM sodium citrate pH 7) as transfer buffer. After blotting the DNA was crosslinked to the membrane with 120 mJ using a UV-Stratalinker (Stratagene).

For the generation of isotope-labelled probes, templates were amplified via PCR, using the primers P38 and P40 containing the T7 promoter sequence together with the respective primers P37 and P39 and genomic DNA from *Synechocystis* 6803. Transcript probes labelled with [α-^32^P]-UTP (3000 Ci/mmol, 10 mCi/mL) were generated from these templates using the MAXIscript® T7 In Vitro Transcription Kit (Thermo Fisher Scientific).

Hybridization was performed overnight at 52°C in hybridization buffer followed by 10 min wash steps each in wash buffers 1, 2 and 3 (for buffers see northern blot section) at 47°C. The signals were detected with a Phosphor Imaging Screen (Bio-Rad) and the GE Healthcare Typhoon FLA 9500 imaging system.

### Endoribonuclease assays

Purification of 6xHis-tagged RNase E was performed as published (Behler et al., 2018). In short, codon-optimized and TEV site-fused *slr1129* from *Synechocystis* 6803 was expressed under control of an IPTG-inducible promoter on pQE70. Expression was induced with 1 mM IPTG in *E. coli* M15 [pREP4] at an OD_600_ of 0.7 at 22°C for 24 h. The cells were collected by centrifugation and the recombinant protein was column-purified using Ni^2+^-NTA resin (Qiagen). The purified protein was stored at -80°C. Elution fraction 2 was used for subsequent cleavage assays.

*In vitro* transcription of *ssr7036* (consisting of the complete 5’ UTR and the first 100 nt of coding sequence) and asRNA1 substrates for RNase E assays was performed with the MEGAshortscript T7 transcription kit (Thermo Fisher Scientific). Suitable templates were PCR-amplified (primers P41 to P44) and thereby tagged with a T7 promoter. The amplified and purified fragments were used for *in vitro* transcription according to the manufacturer’s specifications. *In vitro* transcripts were gel purified using 8 M urea 10% PAA and the ZR small-RNA PAGE Recovery Kit (Zymo Research). The *in vitro* transcribed RNAs carry three additional G nucleotides at their 5΄ ends originating from the T7 promoter. To generate suitable substrates for RNase E, the transcripts were dephosphorylated by RNA polyphosphatase (RPP, Lucigen) generating a 5’monophosphate. For RPP treatment 5 µg gel purified RNA, 20 U RPP, 2 µL RPP buffer and 0.5 µL RiboLock (Thermo Fisher Scientific) were incubated in a total volume of 20 µL for 30 min at 37°C and subsequently column-purified using RNA Clean and concentrator 5 (Zymo Research).

Cleavage reactions were performed in a volume of 10 μL in RNase E cleavage buffer (Behler et al., 2018). Transcripts (0.4 µM) and buffer were premixed, incubated for 5 min at 65°C, cooled down to room temperature, then 2 µL of RNase E or dilutions as indicated were added. As negative control, 2 µL elution buffer 1 were added. Reactions were incubated for 15 min at 30°C, stopped by the addition of 2× RNA loading dye (95% formamide, 0.025% SDS, 0.025% bromophenol blue, 0.025% xylene cyanol FF, 0.5 mM EDTA, pH 8). Half of the reactions were loaded onto denaturing 8 M urea 10% PAA gels. RiboRuler low range RNA ladder (Thermo Fisher Scientific) and low range ssRNA ladder (New England Biolabs) were used as size markers. RNA was visualized with SYBR® Gold Nucleic Acid Gel Stain 1:10,000 diluted in 0.5x TBE. Signals were detected with the biomolecular imager Typhoon FLA 9500 (GE Healthcare). Excitation of 473 nm, emission filter long pass blue ≥ 510 nm and a photomultiplier value of 400-550 V were used.

### Analysis of the genomic structure of *slr7037* knockout strains

Genomic DNA was extracted from cyanobacterial cells as described (Kanesaki et al., 2012). The purity of the genomic DNA extract was checked in an Agilent 2100 bioanalyzer (Agilent Technologies, Palo Alto, Calif.). Since the *Synechocystis* 6803 genome contains many duplicated sequences exceeding 1 kb, such as transposons, we selected a mate-pair sequencing approach, suitable for longer DNA fragments (2 to 10 kb) to analyse the genomic structure. Sequencing libraries were prepared using the Nextera Mate Pair Library Preparation Kit (Illumina). Mate-pair sequencing was carried out for 150 cycles using the Nextseq550 system (Illumina Inc., CA, USA) according to the manufacturer’s specifications. Original sequence reads were deposited in the DRA/SRA database with accession numbers DRX398828 to DRX398831.

The sequencing reads were trimmed using the CLC Genomics Workbench ver. 21.0.3 (Qiagen, Venlo, Netherland) with the following parameters: Phred quality score > 30; removing the terminal 5 nt from both 5′ and 3′ ends; and removing truncated reads <20 nt. The junction adapter sequences originating from the library construction were removed to avoid obstructions to read mapping according to the instructions. 4,000,000 trimmed reads were randomly extracted and mapped to the reference genome sequence of *Synechocystis* 6803 (accession numbers: NC_020286.1, NC_020287.1, NC_020288.1, NC_020289.1, NC_020290.1, NC_020296.1, NC_020297.1, NC_020298.1) using CLC Genomics Workbench ver. 21.0.3 (QIAGEN) with the following parameters: length fraction: 0.8, similarity fraction: 0.9.

The procedure for investigating the integration of pSYSA into the chromosome or other plasmids is shown in **Figure S2**. First, pairs were selected in which one side of the paired reads mapped to pSYSA. Then, this set was narrowed down to pairs in which the other side mapped to a chromosomal or plasmid locus. The same procedure was performed by switching references, and read pairs common to both subsets were extracted. The first position of each read is plotted in **Figure S2**.

The average read depth was calculated by dividing the number of reads mapped to a specific replicon by the length of this replicon. The ratio of read depths of plasmids to chromosome were used as measure for the plasmid copy number relative to chromosomal copies (**Supplemental Table S1**).

## RESULTS

### The most abundantly transcribed region on *Synechocystis* 6803 plasmid pSYSA consists of an mRNA:antisense RNA locus

We observed previously that RNase E expression level affected pSYSA plasmid copy number (Hoffmann et al., 2021). Because overlapping transcripts play a central role in the copy number control of various plasmid replication systems (Lin-Chao and Cohen, 1991; Brantl et al., 1993; Hiraga et al., 1994; Malmgren et al., 1996, 1997), we searched for a pair of overlapping abundant transcripts on the plasmid pSYSA. The most abundantly transcribed region beyond the three CRISPR systems on pSYSA is indeed a locus transcribed from both strands. It contains the short protein-coding gene *ssr7036* on the forward strand and a partially overlapping asRNA, here called asRNA1, transcribed from the reverse strand (**Figure 1**). The *ssr7036* mRNA originates from a single transcriptional start site (TSS) at position 32,177 on the forward strand. The previous classification of transcripts assigned the mRNA to two separate transcriptional units (TUs), TU7029 and TU7030 (Kopf et al., 2014), with TU7030 encompassing the coding sequence of *ssr7036* and beginning just 5 nt before the start codon. However, analysis of PSS/TSS data from the TIER-seq data set (Hoffmann et al., 2021) revealed that TU7029 and TU7030 derive from a single precursor transcript upon processing by RNase E.

**FIGURE 1.**
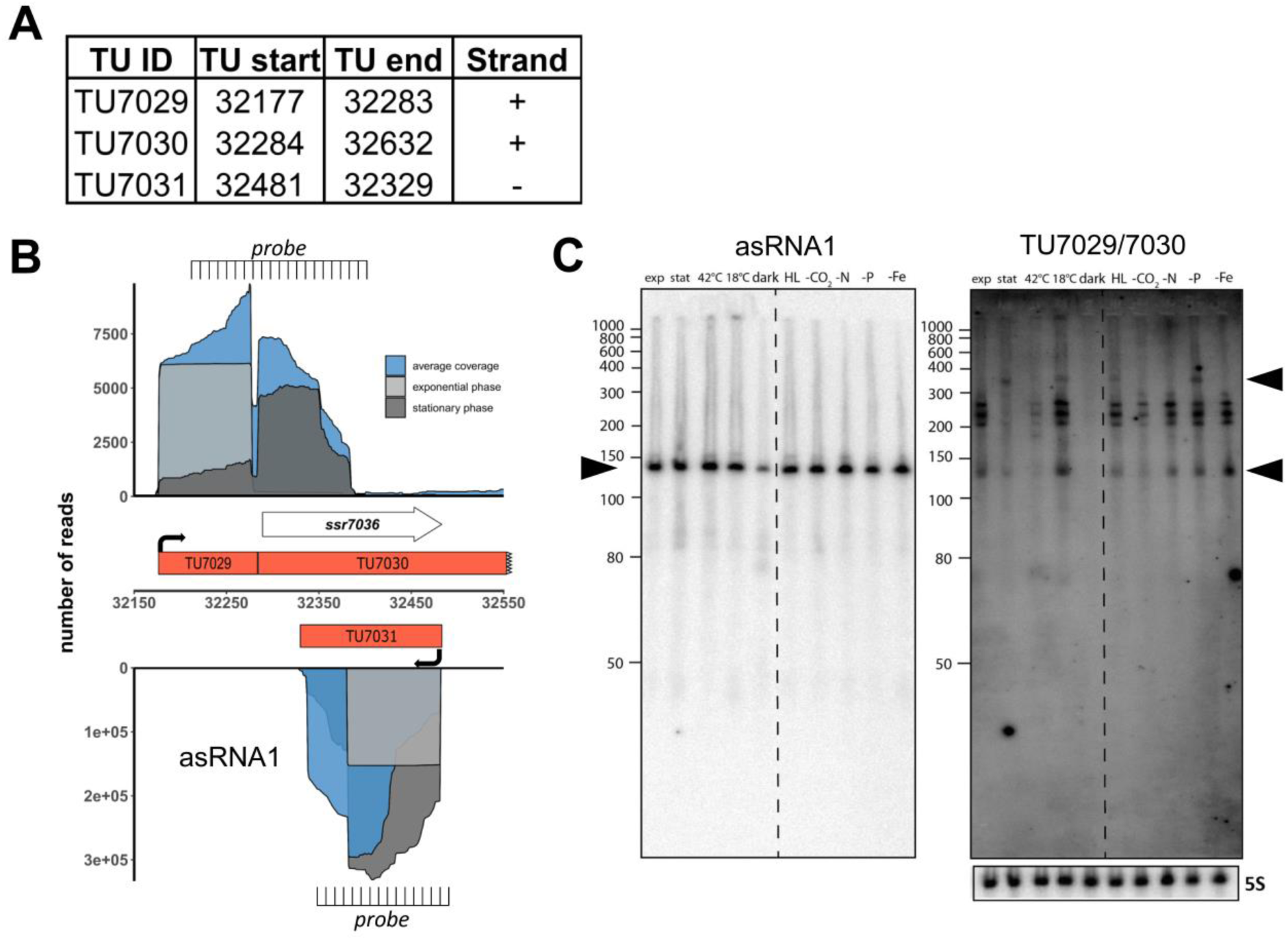
The *ssr7036*/asRNA1 locus on plasmid pSYSA in *Synechocystis* 6803. **(A)** Definition of the transcriptional units TU7029, TU7030 and TU7031 according to previous analyses by dRNA-seq (Kopf et al., 2014). The TU start and end positions are given for the *Synechocystis* 6803 pSYSA sequence available in GenBank under accession number AP004311. **(B)** Location of TUs according to the previous annotation of the transcriptome and genome-wide mapping of transcriptional start sites (Kopf et al., 2014), together with the mapping data from previous dRNA-seq analyses for two conditions, exponential (dark grey) and stationary (light gray) growth phase and the average total coverage (blue). The locations of two probes used in panel C for Northern blot analyses are indicated. **(C)** Northern blot hybridization of total RNA using a ^32^P-labeled transcript probe specific for asRNA1 (TU7031, left) and *ssr7036* (TU7029/TU7030, right panel) subsequently on the same membrane after separation of 12 µg total RNA isolated from cultures grown under ten different growth conditions on a denaturing 10% polyacrylamide gel. The different growth conditions (Kopf et al., 2014) were: exponential phase (exp.), stationary phase (stat.), heat stress for 30 min (42°C), cold stress for 30 min (15°C), darkness for 12 h (dark), high light, 470 µmol photons m^-2^s^-1^ for 30 min (HL), depletion for inorganic carbon, cells were washed 3 times with carbon-free BG-11 and cultivated further for 20 h (-C), nitrogen starvation, cells were washed 3 times with nitrogen-free BG-11 and cultivated further for 12 h (- N), phosphorus starvation, cells were washed 3 times with phosphorus -free BG-11 and cultivated further for 12 h (-P), iron depletion by adding the iron-specific chelator desferrioxamine B and continued cultivation for 24 h (-Fe). The locations of the two probes are indicated in panel B. The asRNA1 and *ssr7036* transcripts accumulate differently under the various growth conditions. Black arrowheads point at the major accumulating transcript form for asRNA1 and in case of *ssr7036,* two transcripts that match approximately the expected lengths for TU7030 and TU7029. A hybridization for the 5S rRNA using the labeled oligonucleotide P62 (Hein et al., 2013) was used for loading control. RiboRuler low range RNA ladder (Thermo Fisher Scientific) and low range ssRNA ladder (New England Biolabs) were used as size markers.

The asRNA1 transcript accumulates to a very substantial level. It originates from a single TSS at position 32,481 on the reverse strand and starts with an adenine which is complementary to the thymidine that is the first nucleotide of the *ssr7036* TGA stop codon. The length of asRNA1 according to differential RNA-seq (dRNA-seq) data (Kopf et al., 2014) is 153 nt (**Figure 1A**). Over its entire length, asRNA1 is complementary to the coding sequence of *ssr7036* (**Figure 1B**). In dRNA-seq data, the asRNA1 level was lowest in the dark, but accumulated to similar levels under all other conditions. In contrast, TU7029 showed widely changing levels under different conditions. The dRNA-seq data are focused on the primary transcript’s 5’ ends, which result from transcriptional initiation (Mitschke et al., 2011). Hence, full-length transcripts can accumulate to a different level than suggested by dRNA-seq. To control for this, we performed Northern blot hybridizations to judge the actual accumulation of asRNA1 and *ssr7036* mRNA. For asRNA1, a minor and a major transcript accumulated slightly above and below 150 nt (**Figure 1C**). This is consistent with the observed dRNA-seq coverage and indicates some heterogeneity at the 3’ end.

For *ssr7036*, signal intensities differed strongly between different conditions. Signals were very low or undetectable for the samples from stationary phase, heat shock at 42°C or darkness, but prominent in samples from exponential growth phase and several other conditions such as cold shock at 18°C, high light or low iron (**Figure 1C**). A particularly long transcript of ∼350 nt was observed in the samples from stationary phase, cold shock, high light and phosphate depletion (upper black arrowhead in **Figure 1C**). Moreover, multiple bands with sizes of ∼140 nt and in the range of 200 to 260 nt (**Figure 1C**) were observed. These multiple bands likely result from processing, because no additional TSS was mapped to this region. One should note that the used probe is complementary to both TU7029 and TU7030.

We conclude that the *ssr7036* mRNA and asRNA1 form a pair of overlapping transcripts originating from pSYSA. The less abundant *ssr7036* mRNA accumulates to different levels in various environmental conditions and appears to be processed at multiple positions out of a precursor transcript spanning TU7029 and TU7030.

### The level of RNase E expression impacts *ssr7036* transcript accumulation

To study possible effects of changing RNase E availability *in vivo*, we used the previously generated strains *rne*(WT) and *rne*(Ts) (Hoffmann et al., 2021). These strains were constructed by introducing the complete *rne-rnhB* locus, including the native promoter and 3’ UTR, on a self-replicating plasmid into *Synechocystis* 6803 WT followed by deletion of the *rne*-*rnhB* genomic locus by homologous recombination (Hoffmann et al., 2021). The strain in which the native, unchanged *rne* gene was introduced was called *rne*(WT). The temperature-sensitive strain *rne*(Ts) was constructed in the same manner, but contains mutations leading to the two amino acid substitutions I65F and V94A in the *rne* gene. A temperature of 39°C was established as leading to RNase E inactivation and causing substantial physiological side effects over time, which were not observed at the permissive temperature of 30°C (Hoffmann et al., 2021). For these strains, we expected the following effects:

- Transcripts accumulating with increasing time at 39°C at a higher level in *rne*(Ts) compared to *rne(*WT) are likely direct targets of RNase E;
- Transcripts accumulating with increasing time at 39°C at a lowered level in *rne*(Ts) indicate RNA species which likely require the enzyme for maturation from a less stable precursor;
- Changed transcript abundance in *rne(*WT) compared to *Synechocystis* 6803 WT result from the higher gene dosage of the plasmid-located *rne-rnhB* operon compared to the chromosomal locus.

We grew cultures of *Synechocystis* 6803 WT, *rne(*WT) and *rne*(Ts) in quadruplicates and subjected them to the non-permissive temperature of 39°C. Total RNA was prepared immediately before the temperature shift (0 h), 1 h and 24 h after. The *ssr7036* transcript and asRNA1 levels were determined by Northern blot hybridization. At 0 h, we observed an increased *ssr7036* signal intensity in *rne*(WT) and *rne*(Ts) compared to *Synechocystis* 6803 WT (**Figure 2A**). This is most likely explained by elevated RNase E levels in *rne(*WT) and *rne*(Ts) compared to WT due to the higher *rne* gene dosage. After 1 h and 24 h at 39°C, the transcript pattern in *rne*(Ts) was markedly changed. A new band appeared in *rne*(Ts) at 1 h while the strongest signals in *Synechocystis* 6803 WT and *rne(*WT) became weakened in *rne*(Ts) at 24 h after the temperature upshift.

**FIGURE 2.**
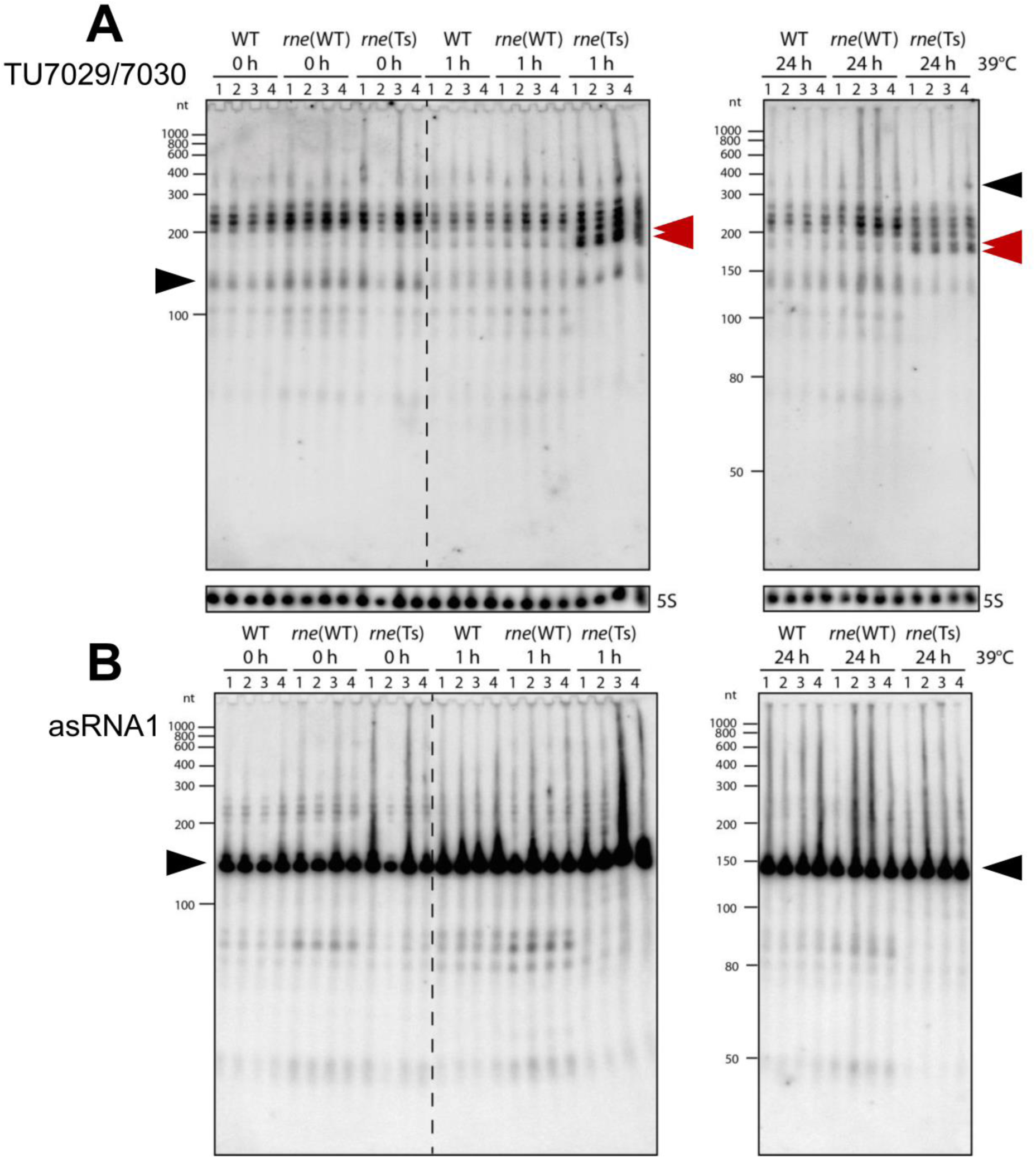
Accumulation of *ssr7036* and asRNA1 in *Synechocystis* 6803 WT. (*rne*(WT) **and a temperature sensitive RNase E variant *rne*(Ts).** Both strains contain an additional plasmid-encoded RNase E gene, resulting in higher RNase E levels compared to WT. Strains were exposed to 39°C to heat-inactivate RNase E in the strain *rne*(Ts) for 0 h, 1 h, or 24 h (four biological replicates). Ten µg total RNA were size-fractionated by denaturing 8 M urea 10% PAGE. **(A)** Hybridization against the TU7029/7030 encompassing *ssr7036.* A double band showing up in *rne*(Ts) at 1 h and 24 h is labeled by two red arrowheads. The same probes were used as in Figure 1C. **(B)** Hybridization against asRNA1. Black arrows as in Figure 1C. Ten µg total RNA were separated per lane and a 5S rRNA hybridization was performed to show equal loading. As molecular mass markers, RiboRuler low range RNA ladder (Thermo Fisher Scientific) and low range ssRNA ladder (New England Biolabs) were used.

For the asRNA1 clear differences were observed in *rne*(Ts) at 1 h and 24 h after the temperature upshift in the accumulation of transcripts just below the 50 nt marker band and in the region between 80 and ∼90 nt (**Figure 2B**). These asRNA1-derived transcripts depended on RNase E activity because the corresponding bands disappeared in *rne*(Ts) with increasing time at the non-permissive temperature.

We conclude that the *in vivo* accumulation of different *ssr7036*-related transcript types changed upon RNase E inactivation.

### Mapping of RNase E cleavage sites *in vivo*

To precisely map the RNase E cleavage sites *in vivo*, we reanalyzed the previously prepared transcriptome-wide datasets of RNase E cleavage sites. After transient inactivation of RNase E (TIER-seq) in the temperature-sensitive strain, *rne*(Ts) (Hoffmann et al., 2021), processing sites with higher read counts in *rne*(WT) compared to *rne*(Ts) indicated RNase E cleavage sites. We identified several RNase-E-dependent processing sites within TU7029, TU7030 and asRNA1 (**Figure 3A**, **Table 3**).

**FIGURE 3.**
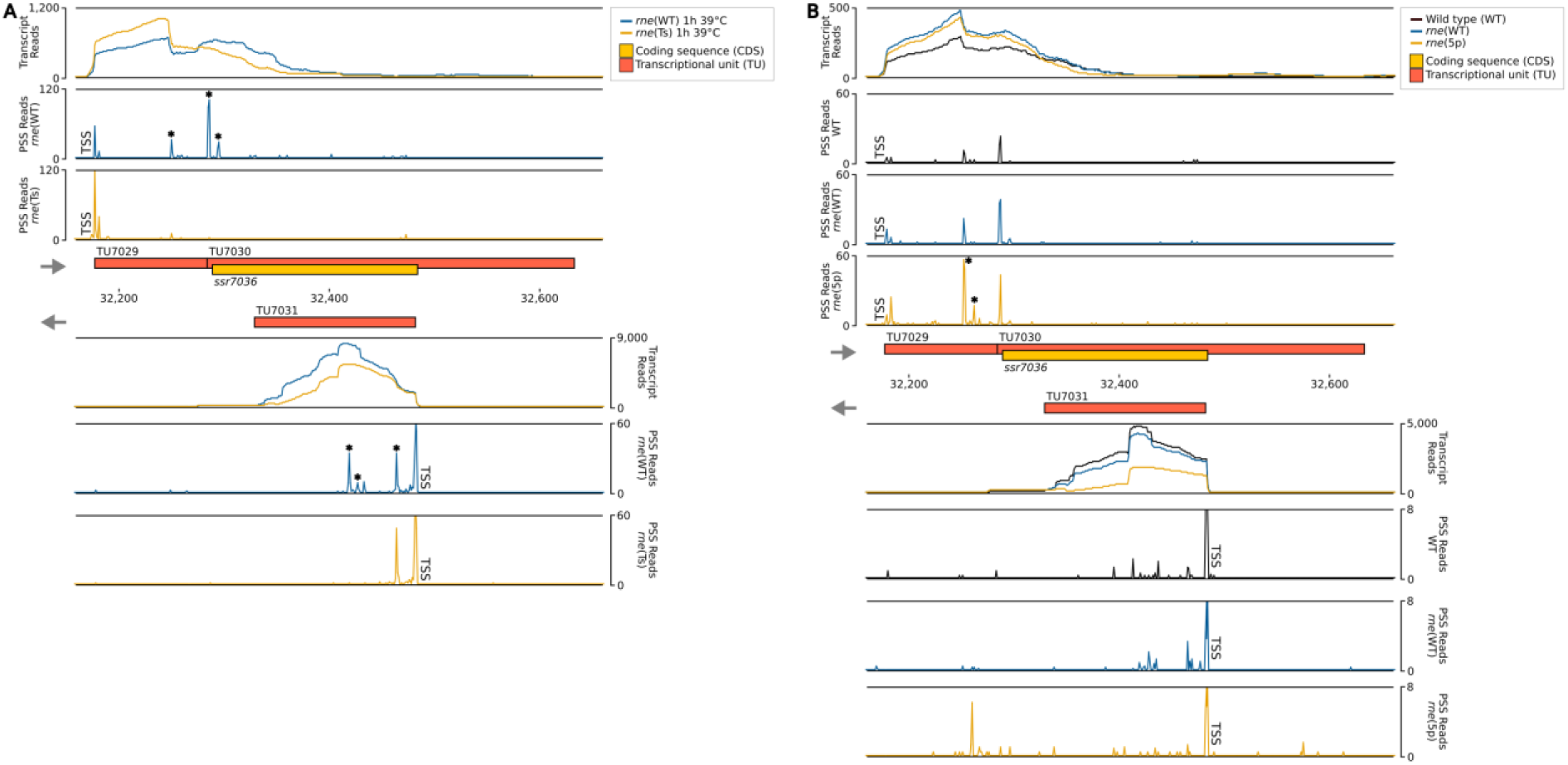
RNase-E-dependent processing events in the *ssr7036*/asRNA1 locus (TU7029, TU7030 and TU7031). **(A)** TIER-seq analysis comparing a strain encoding a temperature-sensitive RNase E variant, *rne*(Ts), to one expressing wild-type RNase E, *rne*(WT), after a heat shock of 1 hour at 39°C. This heat shock inactivates RNase E in *rne*(Ts) and enables the identification of RNase-E-dependent cleavage sites (Hoffmann et al., 2021). **(B)** Comparison of processing sites in wild-type *Synechocystis* 6803 (WT), in strain *rne*(WT) and a strain harbouring 5’-sensing-deficient RNase E, *rne*(5p). For both panels A and B, transcriptome coverage is given on top for the two, respectively three indicated strains. Cleavage sites are displayed in diagrams below by black, blue and orange peaks, representing 5’-monophosphorylated RNA ends (processing sites, PSS) detected in the different strains. 5’-triphosphorylated RNA ends (transcriptional start sites, TSS) may be converted to 5’-monophosphorylated RNA ends *in vivo* or during RNA-seq library preparation. Thus, TSS are partially detected in the PSS signal. Positions which were classified as TSS are indicated by “TSS” next to the respective peaks. Peaks with statistically significantly different read counts between the different strains were indicated by small asterisks next to them (*). Transcriptome coverage and cleavage sites (PSS) represent the average of normalised read counts of the investigated replicates (3 each for *rne*(WT), *rne*(Ts) and WT, 2 for *rne*(5p)).

**Table 3.**
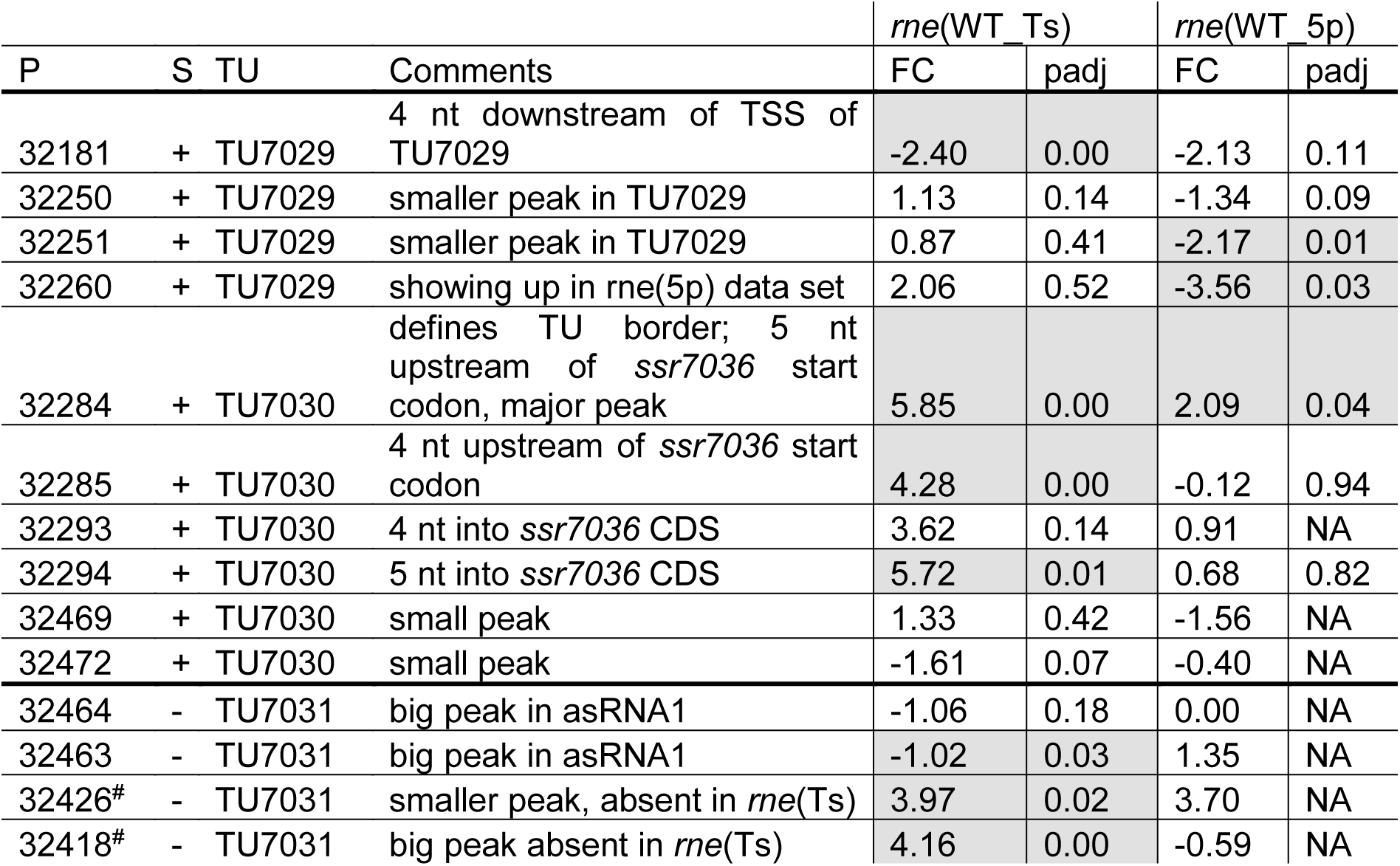
TIER-seq mapped RNase E cleavage sites in the transcriptional units encompassing *ssr7036* and asRNA1. The nt position (P) and strand (S) of the respective site is given on pSYSA, followed by the TU ID and comments. In the last four columns, the comparison for the differences in peak heights are given for *rne*(WT) versus *rne*(Ts) and *rne*(WT) versus *rne*(5p), expressed as log_2_FC (FC) and whether these were significant (if p.adj < 0.05, shaded). Hash symbols (^#^) indicate whether a site matches the results of *in vitro* cleavage analysis in **Figure 4**.

| P | S | TU | Comments | *rne*(WT_Ts) |  | *rne*(WT_5p) |  |
| --- | --- | --- | --- | --- | --- | --- | --- |
|  |  |  |  | FC | padj | FC | padj |
| 32181 | + | TU7029 | 4 nt downstream of TSS of TU7029 | -2.40 | 0.00 | -2.13 | 0.11 |
| 32250 | + | TU7029 | smaller peak in TU7029 | 1.13 | 0.14 | -1.34 | 0.09 |
| 32251 | + | TU7029 | smaller peak in TU7029 | 0.87 | 0.41 | -2.17 | 0.01 |
| 32260 | + | TU7029 | showing up in *rne*(5p) data set | 2.06 | 0.52 | -3.56 | 0.03 |
| 32284 | + | TU7030 | defines TU border; 5 nt upstream of *ssr7036* start codon, major peak | 5.85 | 0.00 | 2.09 | 0.04 |
| 32285 | + | TU7030 | 4 nt upstream of *ssr7036* start codon | 4.28 | 0.00 | -0.12 | 0.94 |
| 32293 | + | TU7030 | 4 nt into *ssr7036* CDS | 3.62 | 0.14 | 0.91 | NA |
| 32294 | + | TU7030 | 5 nt into *ssr7036* CDS | 5.72 | 0.01 | 0.68 | 0.82 |
| 32469 | + | TU7030 | small peak | 1.33 | 0.42 | -1.56 | NA |
| 32472 | + | TU7030 | small peak | -1.61 | 0.07 | -0.40 | NA |
| 32464 | - | TU7031 | big peak in *asRNA1* | -1.06 | 0.18 | 0.00 | NA |
| 32463 | - | TU7031 | big peak in *asRNA1* | -1.02 | 0.03 | 1.35 | NA |
| 32426<sup>#</sup> | - | TU7031 | smaller peak, absent in *rne*(Ts) | 3.97 | 0.02 | 3.70 | NA |
| 32418<sup>#</sup> | - | TU7031 | big peak absent in *rne*(Ts) | 4.16 | 0.00 | -0.59 | NA |

In the second analysed dataset, a strain harbouring a 5’-sensing-deficient RNase E variant, *rne*(5p), was compared to *rne*(WT) and WT. 5’ sensing describes a central mechanism by which RNase E can recognize its targets (Hoffmann et al., in progress). In this analysis, processing sites with higher read counts in *rne*(WT) or *rne*(5p) compared to *Synechocystis* WT indicated cleavage sites of the RNase E native enzyme or the 5’-sensing-deficient enzyme, respectively. Again, several RNase-E-dependent processing sites were identified (**Figure 3B**, **Table 3**).

Altogether, ten PSS were identified within TU7029 and TU7030. Three of these were located at adjacent nucleotide positions and showed a similar response to the manipulation of RNase E activity. The cleavage sites located at position 32,284/32,285, at the border between TU7029 and TU7030, and at position 32,293/32,294, 4/5 nt into the *ssr7036* coding sequence (the reading frame including stop codon extends from position 32,289 to 32,483) (**Figure 3A**) were significantly more prominent in *rne*(WT) compared to *rne*(Ts) (**Table 3**). Therefore, these PSS represent major RNase E processing sites. It is noteworthy that these sites correspond to a pronounced shift in the read coverage. While TU7029 accumulated to a slightly lower level in *rne*(WT) compared to *rne*(Ts), this difference was inversed at the position of these PSS, and TU7030 read coverage was slightly higher in *rne*(WT) than in *rne*(Ts) (**Figure 3A**, upper panel). We conclude that processing by RNase E contributes to a stabilization of TU7029 and a destabilization of TU7030.

Further peaks in TU7029 and TU7030 accumulated at positions 32,250/32,251 and 32,260 in *rne*(5p) compared to *rne*(WT), indicating that a further processing of the respective RNA fragments is dependent on 5’ sensing (**Figure 3B**). No further strong processing sites were detectable within the *ssr7036* coding sequence, except the sites 4/5 nt into the coding sequence. This is likely due to duplex formation with asRNA1, shielding this region from attacks by the single-strand-specific RNase E.

Four processing sites were detected in asRNA1. We noticed that the cleavage site at 32,463/32,464, 18/19 nt downstream of the TSS of asRNA1, was more prominent in *rne*(Ts) than in *rne*(WT) (**Table 3**). Hence, the respective RNA fragments were stabilized due to the declining RNase E activity in *rne*(Ts) after 1 h at the non-permissive temperature. This indicates that this cleavage likely is performed by another RNase, but that the resulting RNA fragment would normally be further processed by RNase E. To the contrary, another two cleavage sites, located at 32,426 and 32,418, accumulated significantly more strongly in *rne*(WT) compared to *rne*(Ts), indicating these were *bona fide* RNase E PSS (**Table 3**).

### *In vitro* analysis of RNase E cleavage sites within the *ssr7036* precursor and asRNA1

To confirm direct RNase E cleavages within the *ssr7036* precursor and asRNA1, *in vitro* cleavage assays were performed. According to the dRNA-seq data set, some asRNA1 transcripts started at the positions just upstream of the main TSS at position 32,481 (**Figure 1A**, **Table 3**). Hence, we included the two preceding nucleotides (positions 32,482 and 32,483 on the reverse strand) in the 153 nt long full-length asRNA1, which was used as a substrate. For the *ssr7036* precursor transcript, we selected a substrate encompassing the entire TU7029 and the first 105 nt of TU7030. Additionally, both substrates contained three additional Gs at their 5’ ends left from the initiation of transcription at the T7 RNA polymerase promoter, yielding transcripts of 158 nt (asRNA1) and 218 nt (*ssr7036*).

The incubation with recombinant RNase E yielded three major fragments for asRNA1 that were not further degraded when increasing the amount of enzyme (**Figure 4A**). Their sizes match the expected lengths of 98, 90 and 60 nt, in case cleavage occurred at sites corresponding to positions 32,426 (yielding the 60 and the 98 nt fragments) and 32,418. The latter cleavage would yield the 90 nt fragment by cleaving off 8 nt from the 98 nt fragment. This interpretation is consistent with the *in vivo* cleavage site mapping, because the processing sites at 32,426 and 32,418 declined in the temperature-sensitive mutant after 1 h at 39°C (**Figure 3**). In contrast, the other site at 32,463/32,464 was likely caused by a different enzyme *in vivo*, as we did not observe fragments matching these positions in the *in vitro* assays (**Figure 4A**).

**FIGURE 4.**
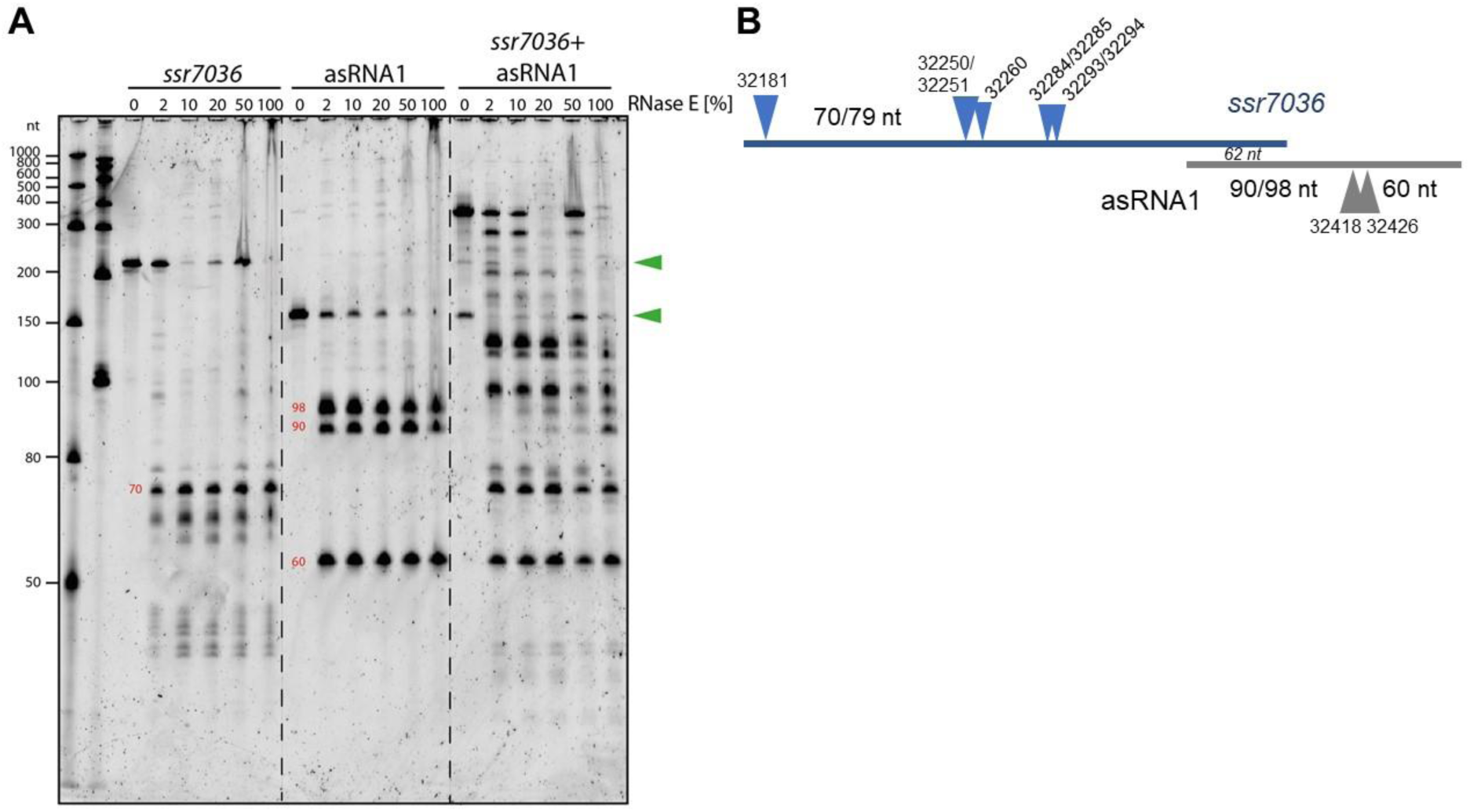
*In vitro* cleavage assays with purified RNase E and potential RNA substrates. **(A)** 5’monophosphorylated *in vitro* transcripts of *ssr7036*, asRNA1 or both were mixed with cleavage buffer, incubated for 5 min at 65°C, cooled down at room temperature and then mixed with 2 µL of recombinant RNase E. Different dilutions of RNase E were used ranging from 1:50 (= 2%) up to undiluted (100%), as indicated. Reactions were incubated for 15 min at 30°C. Half of the reactions were size-fractionated by denaturing 8 M urea 10% PAGE. RiboRuler low range RNA ladder (Thermo Fisher Scientific) and low range ssRNA ladder (New England Biolabs) were used as size markers. RNA was visualized with SYBR® Gold Nucleic Acid Gel Stain. The full-length transcripts are indicated by green arrowheads. The sizes of putative cleavage fragments are given in red numbers. **(B)** Scheme for the cleavage sites and major accumulating fragments in panel A or mentioned in the text.

RNase E processed the *ssr7036* precursor transcript into several lowly abundant fragments and one prominent fragment of approximately 70 nt (**Figure 4A**). When both *in vitro* transcripts were mixed in a 1:1 ratio and incubated with RNase E, larger cleavage products were observed than in the single digests. This indicates the formation of stable RNA duplexes between both molecules. While the band pattern differed quite strongly between the cleavage assays of individual substrates and their mixture, the major 70 nt fragment of the *ssr7036* precursor transcript and the 60 nt asRNA1 fragment did not change. This suggests that these two fragments did not form duplexes in this assay.

The overlapping region between the two substrates was 62 nt (**Figure 4B**). Thus, other signals which did not appear in the cleavage assay with single substrates were also protected by RNA-RNA interactions.

In summary, we could verify several RNase E cleavage sites in both transcripts *in vitro.* The resulting fragments involved the coding region of *ssr7036* and asRNA1 and originated from regions which were not protected by RNA:RNA interactions. The cleavage site at 32250/32251 and the two RNase E sites flanking the start codon of *ssr7036* (32284/32285 and 32293/32294 in **Table 3** were not protected by interaction with the asRNA1.

### Expression of *ssr7036* in *trans* leads to a higher ratio of pSYSA to chromosome

To study a possible effect of *ssr7036* on the pSYSA copy number, we engineered plasmid VIII.17 on basis of the vector pVZ322s in which *ssr7036* was ectopically expressed under control of a rhamnose-inducible promoter (P_rha_). In this construct, the asRNA1 promoter elements are not included and hence asRNA1 is not transcribed from the plasmid. In parallel, a second plasmid, VIII.58, was constructed in which the second codon in *ssr7036* (GAA encoding glutamate) was converted into an UAA stop codon. Except of this single nucleotide difference, the plasmids VIII.17 and VIII.58 were identical.

Southern blot analyses were performed with three independent clones each of the *ssr7036* overexpressor (VIII.17) and the VIII.58 strain containing the *ssr7036* nonsense mutation on the plasmid (VIII.58). As controls we used three technical replicates of the WT and three biological replicates each of strains carrying either the empty vector pVZ322s (plasmid VI.16) or a plasmid in which a *sfGFP* gene was inserted instead of *ssr7036* (VIII.18; see **Table 1** for an overview of all constructs). Prior to DNA preparation, all cultures were divided into two aliquots. One of these aliquots was cultivated in the presence of 3.3 mM rhamnose for 24 h to trigger transcription from the P_rha_ promoter while the other aliquot was not. To analyze the copy number of the pSYSA plasmid relative to the number of chromosomal copies, Southern blot hybridization was performed using two probes. One probe was directed against the unrelated CRISPR3 locus on pSYSA (Scholz et al., 2013), while the other was directed against the *cmpA* locus on the chromosome (gene *slr0040* encoding a bicarbonate binding protein).

The single bands observed for either probe are consistent with the expected HindIII restriction fragment sizes (9.6 kb for *cmpA*; 2.55 kb for CRISPR3) (**Figure 5A**). Moreover, both mutant strains, VIII.17 as well as VIII.58, showed a stronger signal for pSYSA (CRISPR3 probe) relative to the chromosomal signal (*cmpA* probe) compared to the WT and the two other controls. This change in signal intensity was not caused by the presence of the additional plasmid in the cells, because the VI.16 and VIII.18 controls contained the same vector backbone.

**FIGURE 5.**
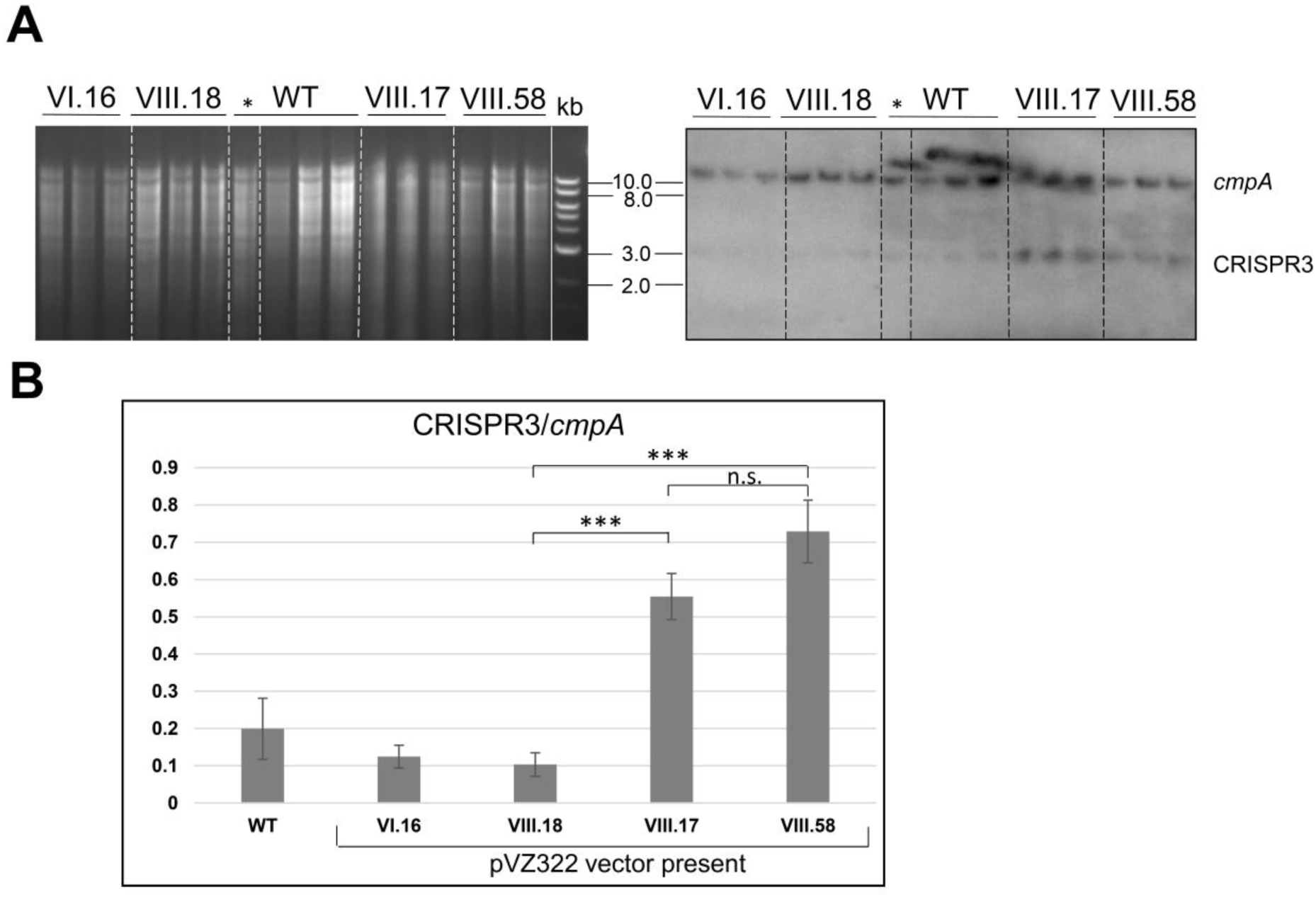
Ectopic overexpression of *ssr7036* leads to enhanced levels of pSYSA. **(A)** Southern blot hybridization against *cmpA* and CRISPR3 (right) and the corresponding gel (left). Four µg HindIII-digested DNA were loaded per lane. Expression was induced by 3.3 mM rhamnose for 24 h except for the one clone of the WT marked with *. WT: wild type DNA; VIII.17: *Synechocystis* 6803 + expressing functional *ssr7036* gene; VIII.58: *Synechocystis* 6803 + expressing *ssr7036* with premature stop codon; VI.16: *Synechocystis* 6803 + pVZ322s control; VIII.18: *Synechocystis* 6803 + GFP control. See **Table 1** for further details of the used plasmids and strains. Three independent clones were used for each plasmid as biological replicates. An 0.8% agarose gel was used for gel electrophoresis. **(B)** Quantification of the signals. Standard errors from triplicates are shown as error bars. An unpaired two-tailed Student’s t-test was performed (see **Table S2** for details). n.s., not significant difference; *** significant difference with p-value < 0.001.

Next, we quantified the obtained differences in signal intensities by dividing the averaged signal intensities for the CRISPR3 fragments by the averaged signal intensities for the *cmpA* fragments. The pSYSA copy number was significantly increased by factors of 2.75 and 3.2 in the VIII.17 and VIII.58 mutant strains compared to the WT (**Figure 5B**). The signal intensities between the VIII.17 and VIII.58 strains were not significantly different, indicating that translation of *ssr7036* was not required for the higher pSYSA copy number. It seems that *ssr7036* transcription was sufficient for this effect. This result supports our idea that the ratio between the *ssr7036* and the asRNA1 transcript amount and their RNase E-mediated cleavage and degradation is important for copy number control of the plasmid pSYSA. In a parallel study, a 3.8-fold increase in the pSYSA copy number was observed in strains overexpressing RNase E and RNase HII (Hoffmann et al., in progress), which matches the results of this work. Hence, we conclude that an elevated *ssr7036* transcript level leads to a higher pSYSA copy number.

### The *ssr7036*/asRNA1 locus together with gene *slr7037* is required for extrachromosomal replication

In the preceding section, we connected the elevated *ssr7036* transcript level to a higher pSYSA copy number. Therefore, we wanted to test if the *ssr7036*/asRNA1 locus could function as an origin of replication. For this purpose, we inserted the *ssr7036*/asRNA1 locus together with the native *ssr7036* 5’ and 3’ UTRs into the plasmid pUC19s yielding plasmid VIII.22 (**Table 1**). The pUC19s origin of replication is not functional in cyanobacteria. Hence, plasmid pUC19s does not replicate in this group of bacteria. Accordingly, transformation of *Synechocystis* 6803 (**Figure S3A**) as well as *Synechococcus* 7942 (**Figure S3B**) was hardly possible with the VIII.22 plasmid, yielding only very few or no colonies, respectively. The experiments were repeated three times, including three technical replicates. We conclude that the *ssr7036*/asRNA1 locus did not function as origin of replication for the modified pUC19s plasmid in *Synechococcus* 7942 and only very rarely in *Synechocystis* 6803. A possible explanation for the observed few colonies for *Synechocystis* 6803 transformation could be a factor present in this organism which is lacking in *Synechococcus* 7942. Indeed, the *ssr7036*/asRNA1 locus is next to the *slr7037* gene, which encodes a 958 amino acid protein with predicted primase and helicase domains (**Figure S4A**), suggesting it might be involved in pSYSA replication. Therefore, we constructed another pUC19s derivative in which the entire contiguous fragment was included, stretching from the *ssr7036*/asRNA1 locus to the end of *slr7037*. The resulting plasmid, VIII.23 (**Table 1**), was used for transformation of *Synechocystis* 6803 and *Synechococcus* 7942 and now colonies were observed for both cyanobacteria (**Figure S3**). We conclude that the entire fragment encompassing the *ssr7036*/asRNA1 locus and the downstream located gene *slr7037* was needed to achieve plasmid replication. However, the numbers of colonies were consistently one to two orders of magnitude higher in *Synechococcus* 7942 than in *Synechocystis* 6803, which likely was due to competition with the native pSYSA plasmid present in the latter. Moreover, we introduced a nonsense mutation into *slr7037* by converting codon 64 into an opal stop codon (**Figure S4B**) yielding plasmid VIII.45. Transformation of VIII.45 yielded no colonies for *Synechococcus* 7942 and a very low number of colonies for *Synechocystis* 6803 (**Figure S4C**). Again, this result can be explained by the pSYSA-encoded native Slr7037 protein in *Synechocystis* 6803 acting in trans.

To test if the plasmid VIII.23 can function as a vector for the expression of cargo genes, we inserted a cassette consisting of the copper inducible *petE* promoter (P*_petE_*) (Zhang et al., 1992) and the *sfGFP* gene encoding the superfolder green fluorescent protein as a reporter, yielding plasmid VIII.44 (**Table 1**). When *Synechocystis* 6803 and *Synechococcus* 7942 were transformed with the plasmid VIII.44, colonies were observed for both strains (**Figure 6**), confirming that the plasmid replicates in these cyanobacteria. Again, the numbers of colonies were one to two orders of magnitudes higher in *Synechococcus* 7942 than in *Synechocystis* 6803. To corroborate the replication of intact plasmids VIII.23 and VIII.44 in these strains, we re-isolated the plasmids from cyanobacteria and successfully transformed *E. coli* with them (**Figure S5**). Moreover, we detected strong sfGFP expression in the transformants of both cyanobacteria, but not in the respective wild types (**Figure 6**). The sfGFP accumulation was inducible by the addition of Cu_2_SO_4_ to a final concentration of 1.25 µM in *Synechocystis* 6803 (**Figure 6A**), while sfGFP was constitutively present in *Synechococcus* 7942, independent from the absence or presence of added Cu_2_SO_4_ (**Figure 6B**). This result is consistent with the presence of a regulatory system that controls the copper-dependent P*_petE_* expression in *Synechocystis* 6803, but is lacking in *Synechococcus* 7942 (García-Cañas et al., 2021).

**FIGURE 6.**
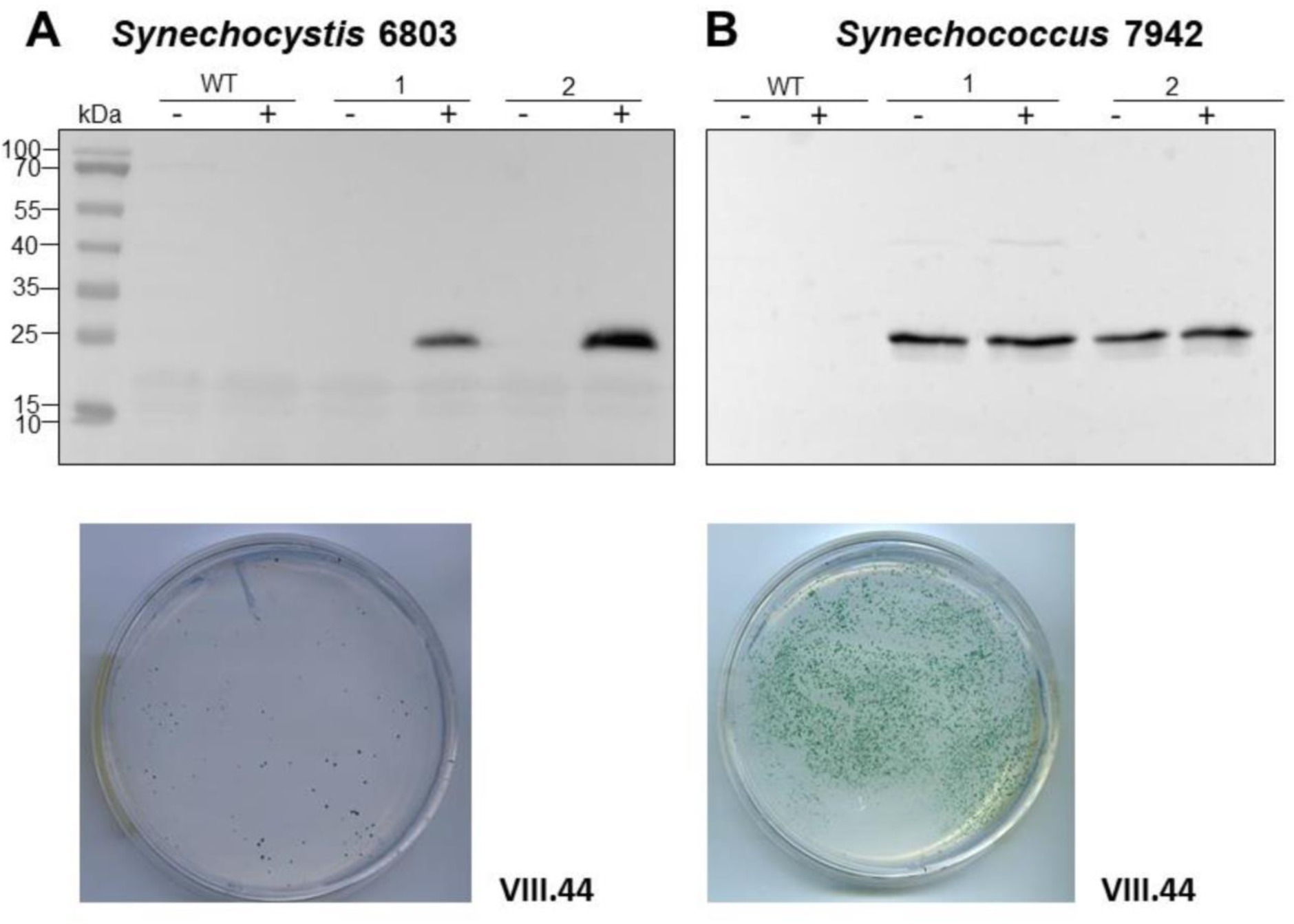
Transfer of plasmid VIII.44 carrying a *sfGFP* reporter gene under control of the P*_petE_* promoter into *Synechocystis* 6803 or *Synechococcus* 7942. **(A)** Expression of sfGFP detected by western blot analysis in *Synechocystis* 6803 after transformation with plasmid VIII.44 which contains the pSYSA origin of replication, i.e. the entire *ssr7036-slr7037* locus. Total protein extracts were prepared from two biological replicates, before and after induction of sfGFP expression by adding Cu_2_SO_4_ to a final concentration of 1.25 µM. The WT was used as negative control. After transformation, selection and re-streaking, the positive clones were grown on plates for 7 weeks and for 1 week in liquid culture before they were collected for western blot analysis. The plate shows an example 2 weeks after transformation with 5 µg of VIII.44 DNA (111 colonies). **(B)** Western blot analysis demonstrating the expression of sfGFP in *Synechococcus* 7942. Total protein extracts were prepared from two biological replicates, before and after adding Cu_2_SO_4_ as in panel (A) but the regulatory system for the control of this promoter does not exist in *Synechococcus* 7942 (García-Cañas et al., 2021). After transformation, selection and re-streaking, the positive clones were cultivated on plates for 4 weeks and for 2 weeks in liquid culture before the analysis. The plate shows a representative example 2 weeks after transformation with 1 µg of VIII.44 DNA (∼1,800 colonies). In both panels (A) and (B), anti-GFP antiserum (Abcam) was used at a dilution of 1:5,000. PageRuler™ Prestained Protein Ladder was used as marker.

Based on these results, we conclude that the *ssr7036*/asRNA1/*slr7037* locus allows the construction of plasmids that can replicate in cyanobacteria and which can be utilized to achieve high expression levels of cargo genes.

### Deletion of *slr7037* leads to integration of the entire pSYSA plasmid into the chromosome or into plasmid pSYSX

Based on the observed dependency of *ssr7036*/asRNA1-primed replication on Slr7037, we wondered if *Synechocystis* 6803 could be cured of the native pSYSA plasmid in the absence of *slr7037*. To answer this question, we generated a deletion strain Δ*slr7037* and examined whether pSYSA was retained in it. Three clones with a complete disruption of the *slr7037* gene (Δ*slr7037*-1,2,3) were selected for further analyses (**Figure S1A** and **B**). Growth of the three clones was comparable to that of the wild type (**Figure S1C**), and sequence analysis showed that they harbored the entire pSYSA sequence (**Table S1**). However, the genomic structure of Δ*slr7037* strains clearly differed from that of the wild type (**Figure 7A**): Read pairs mapped on pSYSA and the chromosome were detected in Δ*slr7037*-1 and -2, indicating that pSYSA was integrated into the chromosome in these clones (**Figure 7B**). In both Δ*slr7037*-1 and -2, pSYSA was integrated into the chromosome via pSYSA genes *slr7104-slr7105*. The fusion sites in the chromosome were *sll0699-sll0700* for Δ*slr7037*-1 and *sll1256-sll1257* for Δ*slr7037*-2. Both *sll0699-sll0700* and *sll1256-sll1257* are homologous genes to *slr7104-slr7105* and encode a transposase. In Δ*slr7037*-2, the number of reads mapped to pSYSA and the chromosome was clearly lower than that observed for the other clones (**Figure 7B**), which seems to be due to a 10-fold increase in the copy number of the small plasmids, pCA2.4, pCB2.4 and pCC5.2 (**Table S1**). There was no evidence for chromosomal integration of pSYSA in the Δ*slr7037*-3 strain. Instead, we observed a fusion between pSYSA and pSYSX (**Figure 7**) via the genes *slr7005* (pSYSA) and *sll6109* (pSYSX). Both genes are identical in sequence and encode putative site-specific integrases. PCR analysis showed complete integration of pSYSA into the chromosome or pSYSX in these strains (**Figure 7C**). These results indicate that pSYSA was not able to replicate as a separate replicon in the Δ*slr7037* mutants, but that its genes were maintained in the chromosome or pSYSX. Therefore, we suggest that Slr7037 functions as a plasmid-encoded replication initiator protein (Rep) protein and renamed it to CyRepA1 (Cyanobacterial Rep protein A1).

**FIGURE 7.**
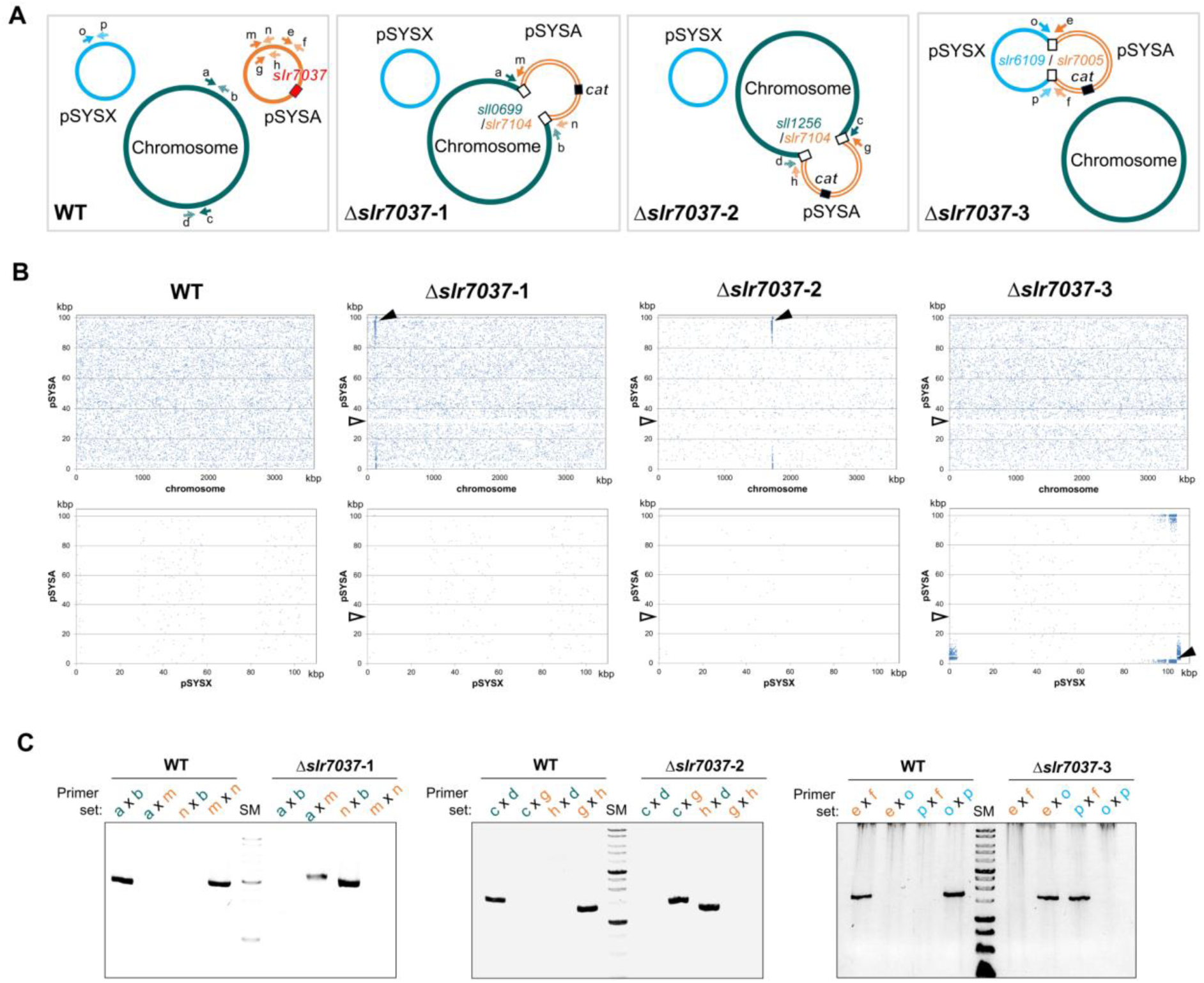
Integration of pSYSA into the chromosome or pSYSX through homologous transposase or integrase genes. **(A)** Genome structure of WT and Δ*slr7037* strains as inferred from sequencing analysis. The positions of *slr7037* in WT, the *cat* gene in Δ*slr7037* mutants and fusion positions between pSYSA and chromosome or pSYSX are shown by filled or open boxes, respectively. The pSYSA plasmid is indicated by the orange double lines. **(B)** The location of paired reads mapped to pSYSA and chromosome (Upper panels) or pSYSX (Lower panels). Blue dots indicate the position of the start of paired reads mapped to pSYSA and chromosomes or pSYSX. Black arrow heads indicate the fusion points between pSYSA and chromosome or pSYSX. White arrow heads indicate the location of Δ*slr7037*. **(C)** The fusion sites of pSYSA and chromosome or pSYSX were amplified using genomic DNA extracted from WT and Δ*slr7037* strains as templates with their specific primers shown in panel A. The thick bands in the size marker (SM) lanes are 3 kb and 1 kb, respectively.

## DISCUSSION

### A pSYSA-derived plasmid for engineering of cyanobacteria

Here we provide evidence that the 3,566 nt long DNA fragment (positions 32,057 to 35,622 on plasmid pSYSA), which contains the *ssr7036* and *slr7037* genes, enabled a pUC19 derivative to replicate independently in *Synechocystis* 6803 and *Synechococcus* 7942. Thus, this DNA region contains an origin of replication which is functional in these two cyanobacteria. We assume that this might also be the case in further cyanobacterial strains. We show that the resulting vector VIII.23 can be engineered to express a reporter gene (plasmid VIII.44; **Figure 6**). Hence, it can be used as a shuttle vector between cyanobacteria and *E. coli*. The presence of VIII.23 in *Synechocystis* 6803 and in *Synechococcus* 7942 and of VIII.44 in *Synechococcus* 7942 was verified 8 weeks after transformation by re-transformation of plasmid DNA isolated from the cyanobacteria into *E. coli* (plasmid re-isolation, **Figure S5**). Thus, plasmid VIII.23 was maintained despite a possible plasmid incompatibility with the native pSYSA plasmid in *Synechocystis* 6803 at least for this period of time. For *Synechococcus* 7942, these results unequivocally support the applicability of VIII.23- and VIII.44-derived plasmids as shuttle vectors with *E. coli*.

### The function of RNase E in the pSYSA copy control mechanism

The initial finding leading to this work was the higher pSYSA copy number observed in strains *rne*(WT) and *rne*(5p) compared to *Synechocystis* WT (Hoffmann et al., 2021). These strains are characterized by an increased level of RNase E relative to the WT, either in its native or 5’ monophosphate sensing deficient form. RNase E is an essential enzyme in cyanobacteria with multiple functions in RNA maturation, degradation and in the post-transcriptional regulation of gene expression (Zhang et al., 2022).

The identification of the *ssr7036*/asRNA1 locus on pSYSA, from which two partially overlapping transcripts originate, provides a link between RNase E and pSYSA copy number control. We show by the analysis of RNA-seq data, *in vitro* cleavage assays and Northern blot hybridizations that both transcripts are processed by RNase E at multiple sites (**Figures 2**, **3** and **4**). We observed that the most striking effects on the *ssr7036* transcript were caused by cleavage at positions 32,260 and at the twin cleavage sites ∼5 nt upstream of the start codon and ∼5 nt into the coding region of *ssr7036* (cf. **Table 3**). These processing events were easily identified in transcriptomic data and these data also showed an inverse effect on the upstream (TU7029) and the downstream (TU7030) segments of the precursor transcript: Whereas TU7029 accumulated to a higher level after the inactivation of the temperature-sensitive RNase E variant at 39°C, TU7030 accumulated to a lower level (**Figure 3A**). In contrast, the higher amount of RNase E in *rne*(WT) led to the stabilization of TU7030, leading to a higher level of some processing products. The longest of these TU7030 transcripts finish within the *ssr7036-slr7037* intergenic spacer (**Figure 1A** and **3A,B, upper panels**), where pSYSA replication likely is primed.

### Comparison of pSYSA replication control with ColE1 and other plasmid systems

Our data suggest that pSYSA is subject to theta plasmid replication, in which the leading strand of a circular plasmid is initiated at a predetermined site (reviewed by (Lilly and Camps, 2015)). In some instances of theta plasmid replication, melting of the DNA double strand depends on transcription, while plasmid-encoded trans-acting Rep proteins can also play a role. Our results show that Slr7037 is essential for maintenance of pSYSA as an autonomous replicon, defining it as a Rep protein. Phylogenetic analysis revealed that homologs of Slr7037 are widely conserved among cyanobacteria (Sakamaki et al., 2022), thus we propose to call this protein CyRepA1. *Synechocystis* 6803 encodes a second Rep protein, ORF B on the small plasmid pCC5.2 (55% similar and 38% identical amino acids). pCC5.2 replicates through the rolling-circle mechanism (Xu and McFadden, 1997). Since ORF B does also have replication activity in cyanobacterial cells, we recently proposed to rename it to CyRepA2 (Sakamaki et al., 2022). Intriguingly, even though CyRepA1 and CyRepA2 are involved in different modes of replication, their protein sequences and structures are very similar.

The copy number of theta plasmids is controlled at the initiation of replication and frequently involves RNAs in antisense orientation to an RNA that is essential for replication, e.g. by acting as a primer for the initiation of replication. Among the best studied examples for this type of copy number control are the ColE1-like origins of replication, which have been fundamental for the development of various expression vectors. In the *E. coli* ColE1 system, RNAI and RNAII, two overlapping, non-coding RNAs play crucial roles. RNAII is the RNA that primes plasmid DNA replication. The level of RNAII is regulated via base-pairing to RNAI, a second, shorter RNA that is much more abundant and sometimes divergently regulated. The TSSs from which RNAI and RNAII originate are 108 bp apart. RNAI has a typical structure consisting of three extended stem-loop elements (**Figure 8**). The level of RNAI is controlled by RNase E that cleaves a pentanucleotide from its 5’ end, destabilizing it effectively (Morita and Oka, 1979; Lin-Chao and Cohen, 1991). In *Synechocystis* 6803, the TSSs of asRNA1 and *ssr7036* are more distantly localized (304 nt). Judged by the number of reads in prior RNA-seq analyses, asRNA1 is on average more than 100 times more abundant than *ssr7036* (**Figure 1**). Compared to RNAI, the asRNA1 is longer (∼150 nt compared to 108 nt), but the predicted secondary structure of asRNA1 resembles RNAI by the presence of three stem-loops (**Figure 8**). Therefore, it is possible that it uses also a similar mechanism to contact its cognate partner molecule, the coding section of the *ssr7036* mRNA, via kissing complexes, i.e. base pairing between complementary sequences at the respective loop sections as described for the RNAI:RNAII interaction (Eguchi et al., 1991).

**FIGURE 8.**
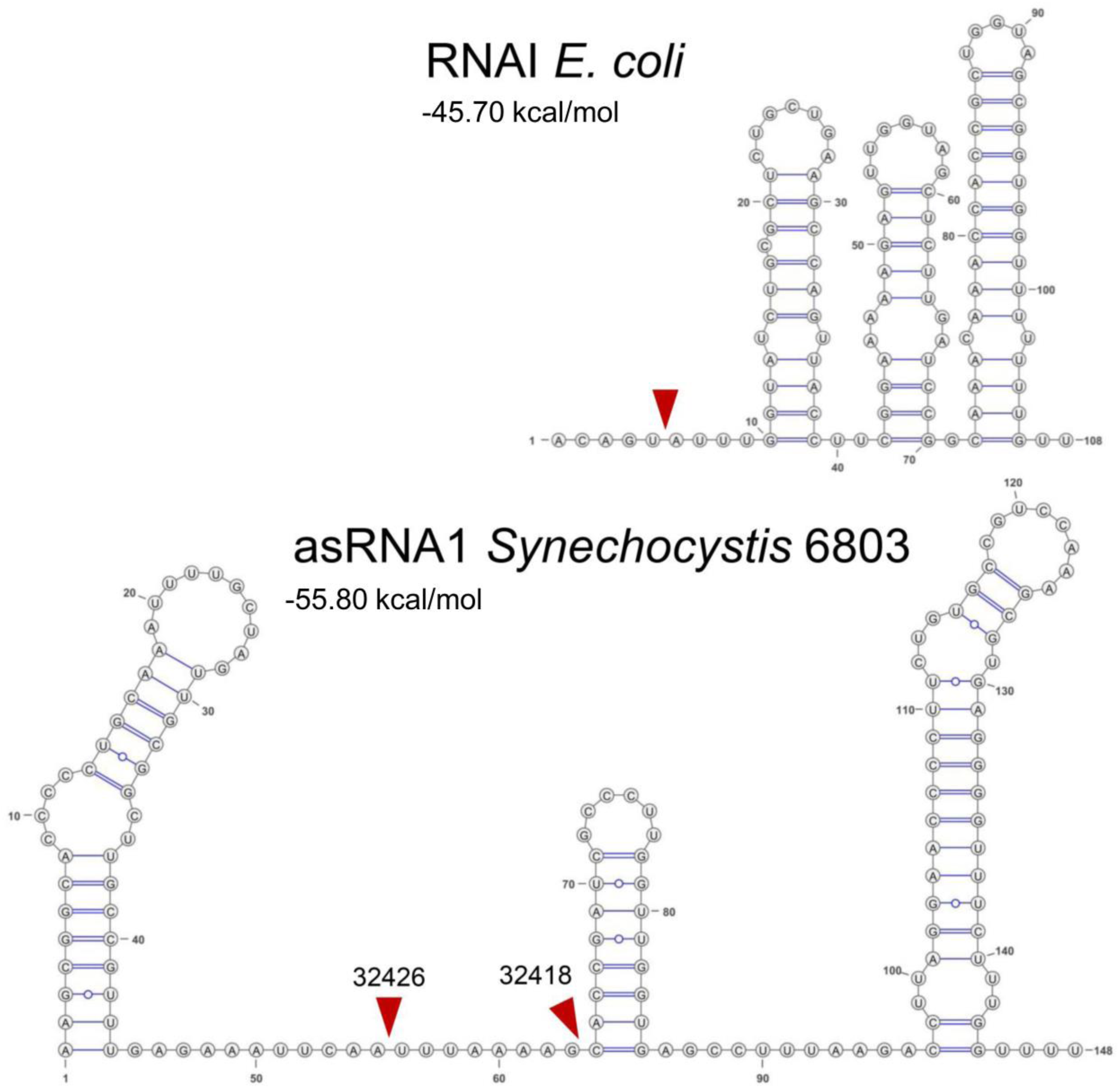
Secondary structures and mapped RNase E cleavage sites in the ColE1 RNAI and the pSYSA asRNA1. Both transcripts are with 108 and ∼150 nt relatively small non-coding RNAs and both overlap a substantial section of another transcript, RNAI in case of the ColE1 replicon and the *ssr7036* mRNA in case of the pSYSA plasmid. Their secondary structures are shown here together with mapped RNase E cleavage sites (read arrowheads) according to this work (asRNA1) or previous analyses for RNAI (Morita and Oka, 1979; Lin-Chao and Cohen, 1991). Sequences of *E. coli* RNAI can be found in the RFAM database (Kalvari et al., 2018) under the accession number RF00106 and the here shown sequence was taken from the European Nucleotide Archive (https://www.ebi.ac.uk/ena), accession no. S42973. The RNA secondary structures were predicted using RNAfold as part of the ViennaRNA Web Services (Gruber et al., 2015) with default parameters, in case of RNAI manually adapted according to published structural models (Morita and Oka, 1979; Lin-Chao and Cohen, 1991) and visualized with VARNA version 3.93 (Darty et al., 2009). The minimum free energy values are indicated in kcal/mol as predicted by RNAfold.

However, there are also major differences between the pSYSA and the ColE1 systems. One of these is the presence of at least two RNase E cleavage sites in asRNA1, which are not close to its 5’ end as in the ColE1 RNAI but within the single-stranded region of the asRNA1 molecule separating stem-loops 1 and 2 from each other (**Figure 8**). It is likely that these cleavage events destabilize asRNA1. Upon overexpression of RNase E we observed the putative resulting cleavage fragments by Northern blot hybridization. Though the pattern suggested their rapid further turnover (**Figure 2B**). Another difference is the presence of a small protein, called Rop (repressor of primer), respectively Rom (RNA one modulator), a 63 amino acids protein relevant for ColE1-type replication encoded downstream of the origin of replication (Eguchi and Tomizawa, 1990). Acting as an adaptor protein (Helmer-Citterich et al., 1988), Rom/Rop enhances the binding between RNA I and RNA II, thereby increasing the inhibitory activity of RNA I (Tomizawa and Som, 1984). Accordingly, Rom/Rop defective mutants show an increased plasmid copy number (Vieira and Messing, 1982). Similarly, the pSYSA system includes a small protein (Ssr7036) as well. It consists of 64 amino acids, but unlike Rom/Rop, it is not encoded in the vicinity but within one of the interacting RNAs. Moreover, the predicted basic IEP of Ssr7036 differentiates it from the acidic Rom/Rop. However, most importantly, we observed that the ectopic expression of a second *ssr7036* gene copy led to a significantly higher pSYSA copy number compared to the chromosome (**Figure 5**). This effect was also clearly observable if the additional *ssr7036* copy contained a stop codon. Therefore, it was caused by the enhanced transcript level, irrespectively of any hypothetical protein function. The interaction with ribosomes and translation of *ssr7036* RNA may stabilize and protect the RNA from degradation. Therefore, in contrast to Rom/Rop’s inhibitory function on plasmid copy numbers, Ssr7036 have a rather additional positive effect on the pSYSA copy number.

Our model summarizes the roles of the two overlapping small RNAs, RNase E and the CyRepA1 protein in pSYSA replication (**Figure 9**). The *ssr7036* mRNA is transcribed as part of a precursor transcript that is processed by RNase E at two closely spaced major cleavage sites into two transcriptional units, TU7029 and TU7030. Several minor cleavage sites were detected as well. The asRNA1, an abundant transcript that is complementary to the coding sequence of *ssr7036,* accumulates in parallel. Therefore, the protein Ssr7036 can only be produced from mRNA species that are not truncated by RNase E and if translation is not hampered due to interaction with asRNA1. Such *ssr7036* transcripts can extend to a position in the *ssr7036-slr7037* intergenic spacer, where replication likely is primed. The downstream gene *slr7037* encodes a protein with predicted helicase and primase domains, typical for a Rep protein involved in plasmid replication. Indeed, our data suggest that Slr7037 is essential for pSYSA replication. However, the genetic information on pSYSA was not lost in the Δ*slr7037* deletion strains because the entire pSYSA plasmid recombined into the chromosome or the pSYSX plasmid. This is consistent with the presence of at least seven different toxin-antitoxin systems on pSYSA (Kopfmann and Hess, 2013), which should mediate a strong post-segregational killing effect if the genetic information would be lost.

**FIGURE 9.**
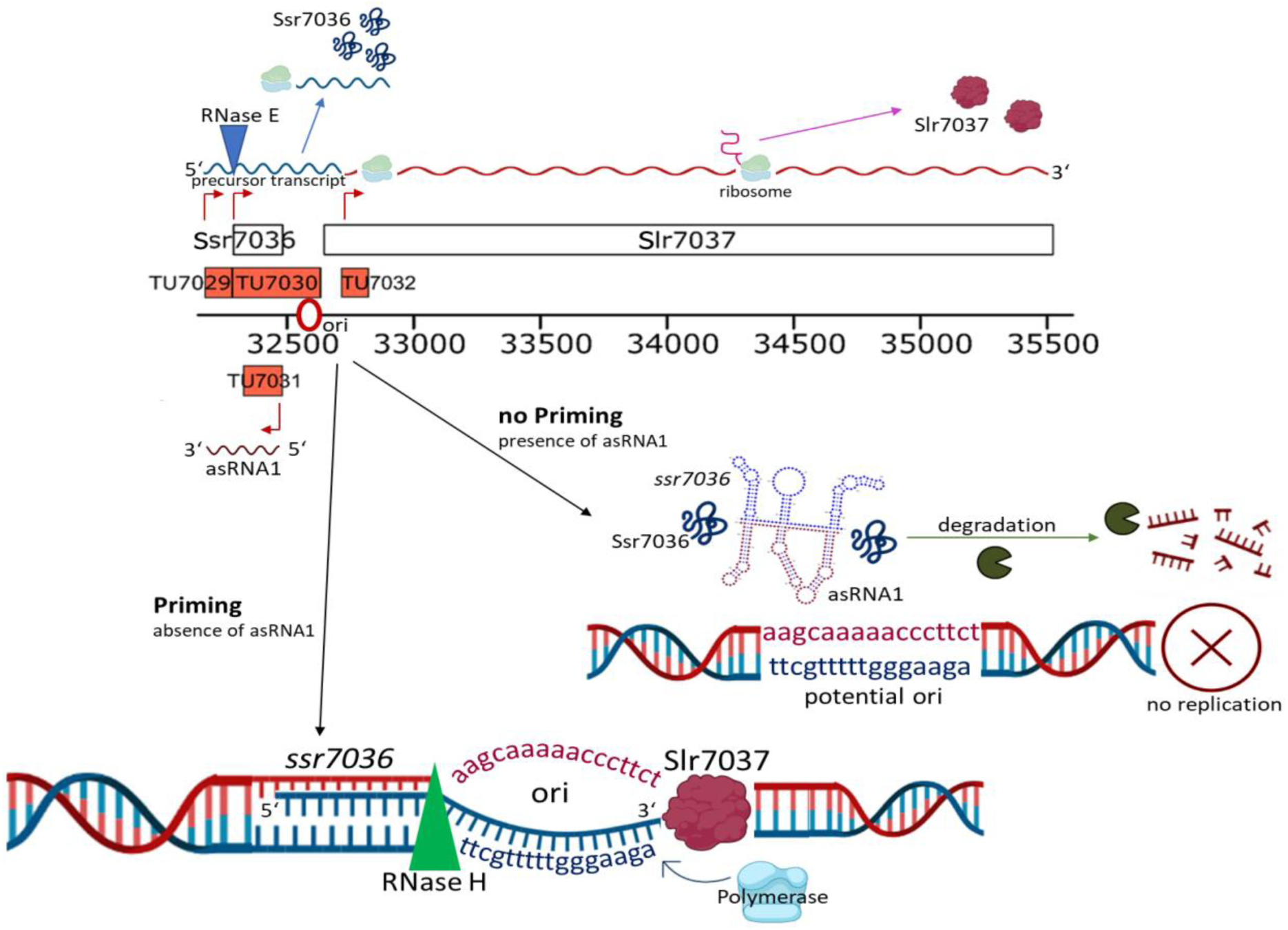
The roles of overlapping small RNAs, RNase E and the Slr7037 Rep protein in pSYSA replication. The *ssr7036* mRNA is central in pSYSA replication. It is processed by RNase E into two transcriptional units, TU7029 and TU7030. An abundant transcript, asRNA1, that is complementary to the coding sequence of *ssr7036* accumulates in parallel. Therefore, the protein Ssr7036 can only be produced from mRNA species that are not truncated by RNase E and if translation is not hampered due to interaction with asRNA1. Ssr7036 may be involved in the interaction between the two transcripts but direct evidence is lacking. Those *ssr7036* transcripts that extend beyond the asRNA1 TSS to a position in the *ssr7036-slr7037* intergenic spacer, are likely those that prime pSYSA replication. The downstream located gene *slr7037* encodes the CyRepA1 protein, as defined in this work. The scheme was drawn using elements from the BioRender platform (https://biorender.com/).

The gathered information led to the construction of plasmids suitable as shuttle vectors for the genetic manipulation of another model cyanobacterium, *Synechococcus* 7942. These provide a solid basis for the development of further vectors and the manipulation of additional cyanobacterial strains or species.

## DATA AVAILABILITY

The resequencing datasets produced in this study are available in the DDBJ Sequence Read Archive DRA/SRA database with the accession numbers DRX398828 to DRX398831.

## AUTHOR CONTRIBUTIONS

WRH designed the study. AK and VR constructed plasmid vectors, engineered cyanobacterial lines and did molecular analyses, VR performed RNase E *in vitro* assays, CS analyzed transcriptomic data, UAH and AW analyzed TIER-seq data, TA, MS, KNM and SW constructed *slr7037* deletion strains and analyzed their genomic structure, AK, VR, SW and WRH drafted the manuscript with input from all authors. All authors read and approved the final manuscript.

## ACKNOWLEDGMENTS

This work was supported by the Deutsche Forschungsgemeinschaft (DFG) Research Training Group Me*In*Bio [322977937/GRK2344] to AW and WRH, through the DFG priority program SPP 2002 “Small Proteins in Prokaryotes, an Unexplored World”, grant HE2544/12-2 to WRH, DFG grant STE 1192/4-2 to CS, and the Ministry of Education, Culture, Sports, Science and Technology of Japan (20K05793) to SW. We thank Ingeborg Scholz and Luisa Hemm (both at the University of Freiburg) for providing the RNA samples in **Figure 1C** and plasmid VII.48, respectively, and John Heap (Imperial College London) for an aliquot of pCK355. The mate-pair sequencing was supported by the Cooperative Research Grant of the Genome Research for BioResource, NODAI Genome Research Center, Tokyo University of Agriculture. We acknowledge support by the Open Access Publication Fund of the University of Freiburg.

