## Supplemental Figures for "Regulation of pSYSA defense plasmid copy number in *Synechocystis* through RNase E and a highly transcribed asRNA"

### Supplemental Material

### SUPPLEMENTAL TABLES

**Supplemental Table S1.** Read depth of mate pair sequence reads

**Supplemental Table S2.** Details of the statistical analysis in **Fig. 5B**.

**Table S1** and **Table S2** are provided together as separate Excel file "Supplemental Tables.xlsx**"**.

### SUPPLEMENTARY FIGURES

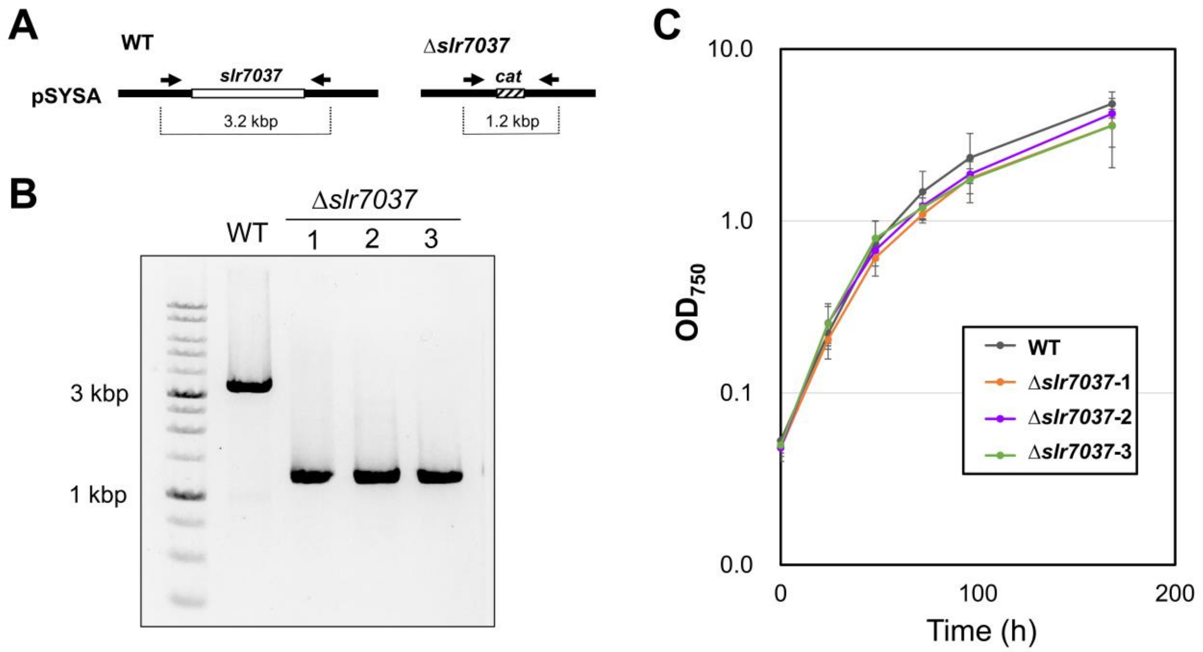


**FIGURE S1 │ Construction of wild type (WT) and *slr7037* knockout strains (Δ*slr7037*) and their growth. (A)** Schematic representation of *slr7037* gene disruption. The chloramphenicol resistance gene cassette (*cat*) was used to replace gene *slr7037* on pSYSA plasmid. **(B)** PCR analysis of the *slr7037* gene locus. The *slr7037* gene locus was PCR-amplified from genomic DNA of WT and three knockout clones with primers represented by arrows in (A). Amplified products were analyzed by 1% agarose electrophoresis. **(C)** Growth curve of WT and Δ*slr7037* strains. Cells cultured on BG-11 plates for one week were harvested and transferred to liquid medium to achieve $OD_{750}$ = 0.05 and cultivated under 50 µmol photons $m^{-2}$ $s^{-1}$ with 2% $CO_2$ bubbling.

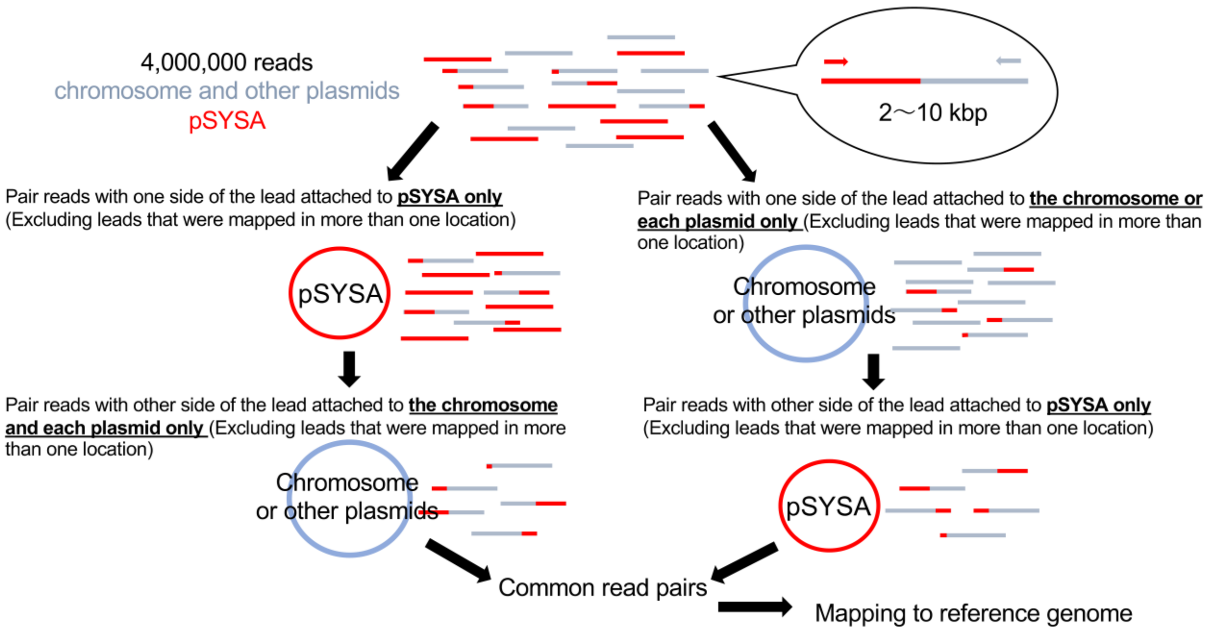


**FIGURE S2 │ Scheme of analysis of mate-pair sequencing reads.** To determine the genomic structure of *Synechocystis* 6803 WT and Δ*slr7037* mutants, 4,000,000 trimmed sequencing reads, obtained by mate-pair sequencing, were analyzed using this scheme. See Materials and Methods for further details.

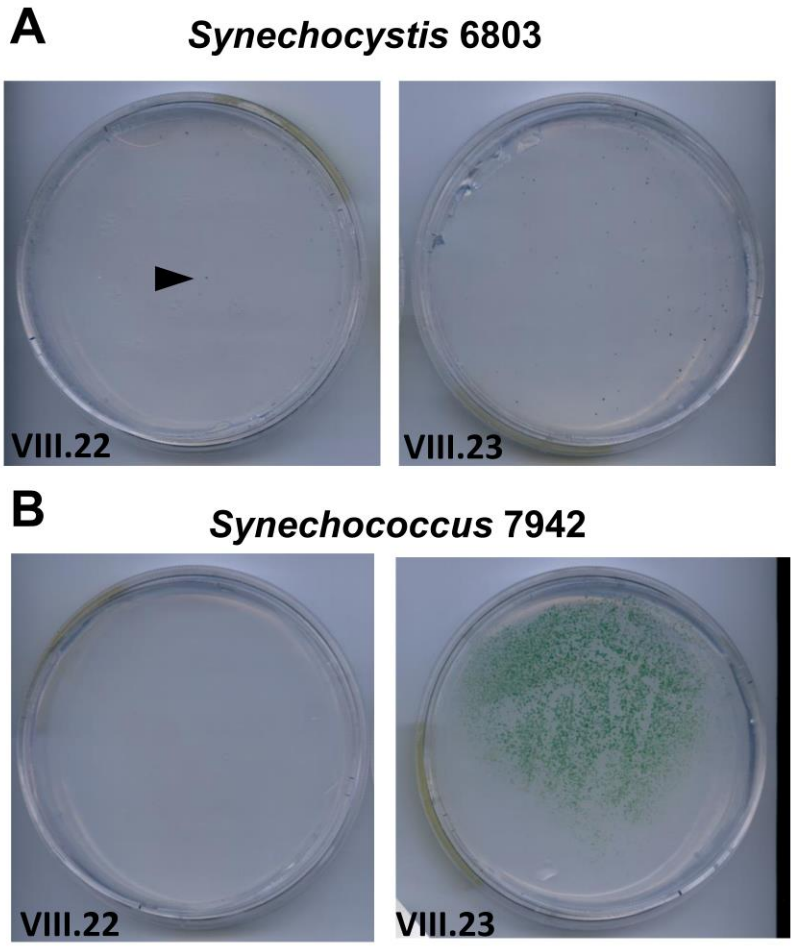


**FIGURE S3 │ Transformation of plasmids containing *ssr7036* (plasmid VIII.22) or the entire *ssr7036-slr7037* locus (plasmid VIII.23) into *Synechocystis* 6803 or *Synechococcus* 7942. (A)** Sometimes single colonies were observed in the transformation of plasmid VIII.22 into *Synechocystis* 6803 (here: 1 colony, arrowhead) containing only *ssr7036* in a pUC19 vector derivative, while consistently some colonies (here: 67) were obtained after transformation with VIII.23 containing the entire *ssr7036-slr7037* locus (**Table 1**). **(B)** High numbers of colonies appeared in *Synechococcus* 7942 upon transformation with VIII.23 containing the *ssr7036-slr7037* locus (here: more than 2,900) but never with plasmid VIII.22. In the shown transformation experiments 5 µg plasmid DNA were added (n=3).

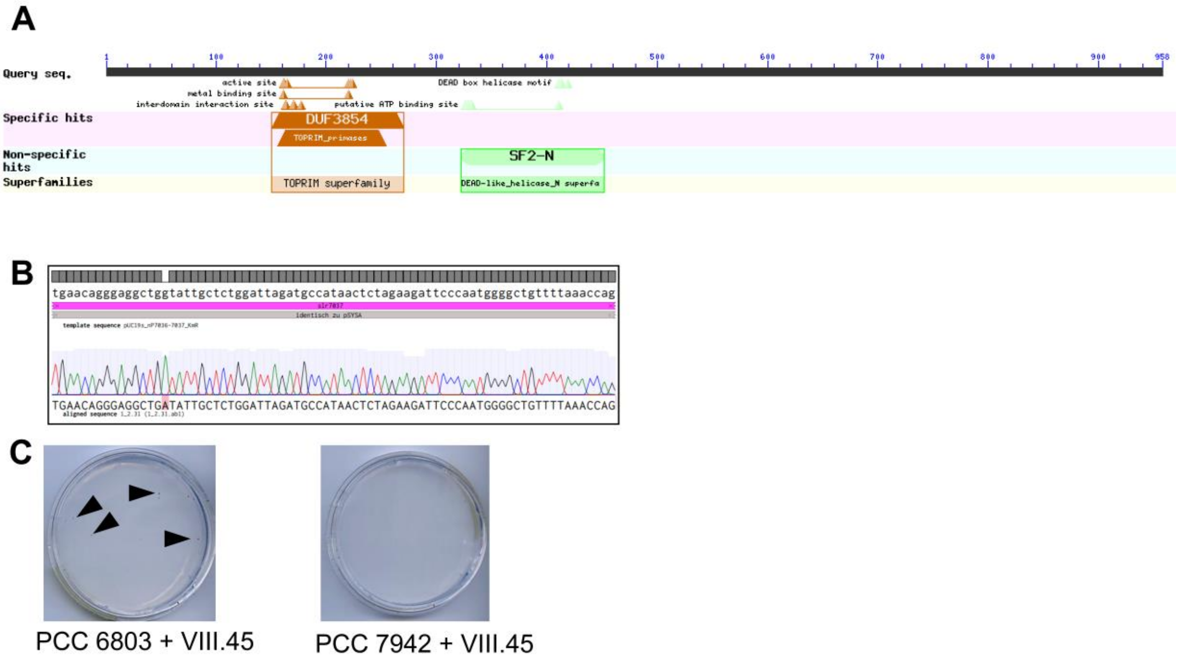


**FIGURE S4 │ Slr7037 as a Rep protein. (A)** Protein domains in Slr7037 predicted by the domain search implemented in the blastP algorithm at https://blast.ncbi.nlm.nih.gov/Blast.cgi?PAGE=Proteins. The topoisomerase-primase (TOPRIM) nucleotidyl transferase/hydrolase domain is found in the active site regions of bacterial DnaG-type primases and their homologs. The TOPRIM domain has two conserved motifs, one of which centers at a conserved glutamate and the other one at two conserved aspartates (DxD). Both of these motifs are present in Slr7037 (E161, D220 and D222). DnaG type primases are often closely associated with DNA helicases in primosome assemblies and indeed there is a DEAD/H-box superfamily 2 helicase domain detectable (here, residues 322 to 452; an extended region predicted by HHpred from residues 318 to 618). **(B)** Mutagenesis of *slr7037*. Codon 64 (TGG) encoding trp (W64) was converted into an *opal* TGA stop codon. The original sequence of VIII.23 (top) was aligned with the respective sequence-verified section in plasmid VIII.45 (bottom). The original G was replaced by A (highlighted in red), leading to the stop codon TGA. **(C)** Transformation of plasmid VIII.45 resulted in small numbers of colonies in *Synechocystis* 6803 (11 in this example, some labelled by arrowheads) and none in *Synechococcus* 7942.

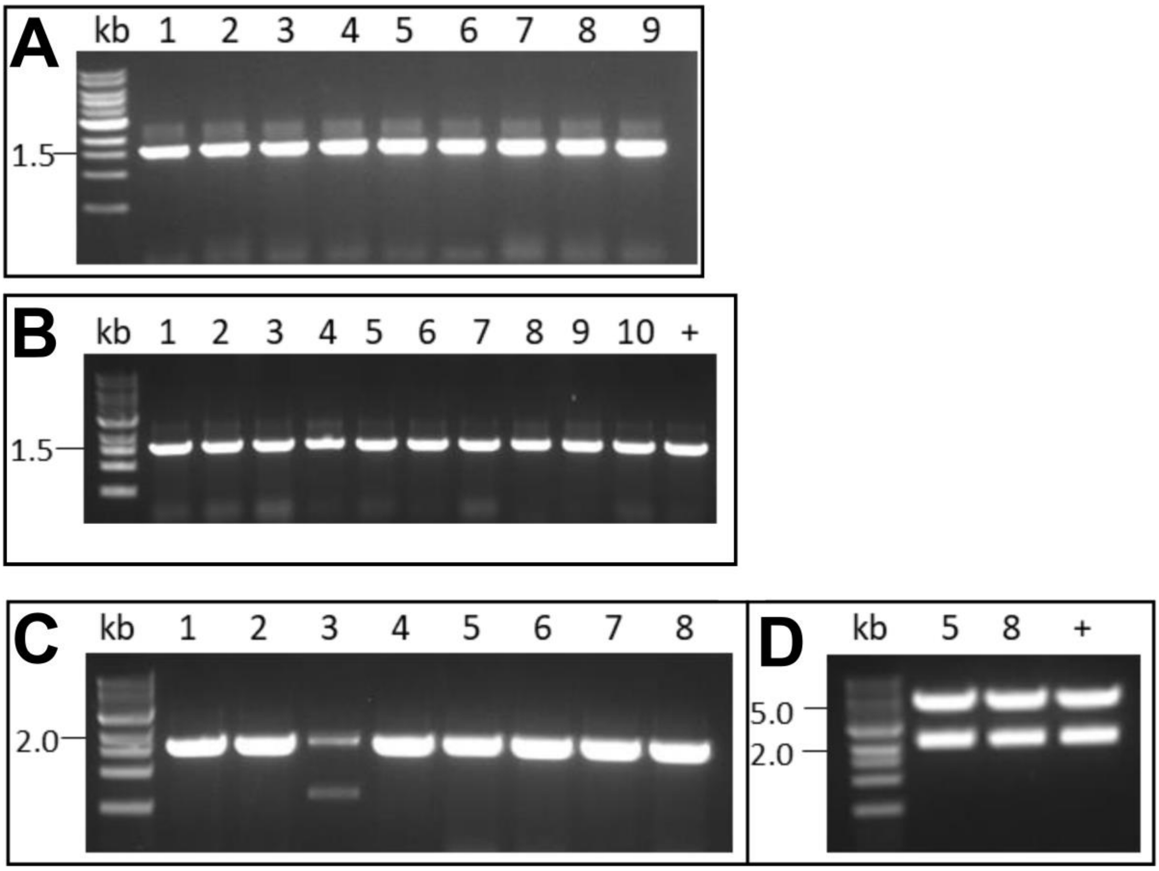


**FIGURE S5 │ Plasmid re-isolation from transformed cyanobacteria and transformation of *E. coli* confirm the presence of intact plasmids. (A)** Total DNA was prepared from cultures going back to two separate *Synechocystis* 6803 colonies that had been transformed with plasmid VIII.23. Competent *E. coli* DH5α cells were transformed with 10 µg total DNA each and selected for the VIII.23 plasmid on 50 µg/mL kanamycin. The presence of the plasmid in 9 randomly selected *E. coli* colonies was confirmed by PCR using primers P57 and P36. A 1.6 kb fragment was expected.

**(B)** Total DNA was prepared from three separate *Synechococcus* 7942 colonies eight weeks after transformation (four weeks on plates and four weeks in liquid culture) with plasmid VIII.23. *E. coli* DH5α was transformed with 10 µg of the isolated total DNA each and selected for the presence of the VIII.23 plasmid on 50 µg/mL kanamycin. The presence of the plasmid in 10 randomly selected *E. coli* colonies was confirmed by PCR using primers P58 and P60. A 1.6 kb fragment was expected. The + symbol indicates a positive control in which 100 pg of plasmid VIII.23 were used for transformation of *E. coli* and one clone was picked.

**(C)** Plasmid VIII.44 re-isolation from *Synechococcus* 7942 eight weeks after transformation as in panel (B), followed by transformation of *E. coli* DH5α and PCR analysis with primers P59 and P61 to verify the presence of the VIII.44 plasmid in *E. coli*. A 1.7 kb fragment was expected and clearly obtained in 7/8 of the analyzed clones.

**(D)** Plasmid DNA was prepared from *E. coli* clones 5 and 8 in panel (C) and a restriction digest performed with the enzyme AatII to verify the presence of the entire plasmid. For the restriction digest with AatII, 2.1 and 5.4 kb long fragments were expected. Bands matching these sizes were obtained. The + symbol refers to a restriction digest of the original plasmid VIII.44 serving as positive control.
